# MLE EEG Toolbox: A Personal AI Laboratory Assistant for Literature-Informed and Reproducible M/EEG Research

**DOI:** 10.64898/2026.11.18.767737

**Authors:** Xiaobo Liu, Wenjian Chen, Bin Wan, Sanwang Wang, Zhiyang Lin, Xihan Zhang, Li Dong, Shiang Hu, Xianwei Guo, Ruifang Cui, Ruiyang Ge

## Abstract

Electrophysiological research increasingly relies on complex analytical workflows, yet integrating methodological evidence, computational execution and reproducibility remains challenging. Here we introduce MLE EEG Toolbox, a framework for building a personal AI laboratory assistant for electroencephalography (EEG) and magnetoencephalography (MEG) research. The toolbox integrates locally deployed AI, natural-language-guided analysis, literature retrieval and methodological comparison, and reproducible computational workflows. A constrained language model translates research instructions into validated, researcher-approved operations, while literature-informed guidance supports parameter selection and protocol reconstruction. Computational provenance enables transparent inspection and replay of analytical workflows. The assistant achieved action-selection accuracies of 85.6% for EEG and 95.5% for MEG, while 142 cross-platform tests demonstrated close numerical agreement. Applications to public EEG and MEG datasets illustrated integrated spectral, network, decoding and source analyses. Running on standard personal computers without external AI APIs or token charges, MLE EEG Toolbox provides an accessible, researcher-controlled environment that connects methodological knowledge with reproducible electrophysiological analysis.

## Introduction

Electroencephalography (EEG) and magnetoencephalography (MEG) provide non-invasive measurements of human brain activity at millisecond resolution, offering complementary perspectives on the temporal and spatial organization of neural dynamics^1–3^. Their temporal precision has made them central to investigations of cognition, brain disorders and large-scale neural interactions ^3,4^. Advances in computational neuroscience have substantially expanded the information that can be extracted from electrophysiological recordings, from oscillatory activity and functional connectivity to transient brain states, complex temporal structure and distributed neural representations^1–3^. Yet the growing sophistication of these approaches has also increased the methodological and computational demands of electrophysiological research.

A central challenge is that EEG and MEG analyses involve a succession of interdependent methodological decisions, each of which can influence the resulting neurophysiological interpretation^5–7^. Choices concerning filtering, artifact removal, temporal segmentation, sensor-space and source-space estimation, feature extraction and statistical inference are often implemented across different computational environments. Established platforms such as EEGLAB, Brainstorm, FieldTrip and MNE have made sophisticated electrophysiological methods widely available, yet integrating diverse analytical procedures into coherent, reproducible workflows can remain technically demanding ^8–11^. Moreover, the computational history linking raw recordings to scientific conclusions is not always preserved in a form that can be independently inspected or reproduced ^12,13^. Consequently, methodological flexibility, although essential for scientific discovery, can also introduce variability that complicates the interpretation and comparison of electrophysiological findings^12,14^.

The emergence of artificial intelligence introduces a new dimension to this challenge. Large language models can interpret natural-language instructions, interact with external tools and translate research objectives into computational actions, potentially reducing the technical expertise required to configure complex analyses ^15,16^. However, generating a plausible analytical workflow is fundamentally different from establishing its methodological validity. Language-model agents may generate executable procedures without reliably satisfying the scientific requirements of a given task, and their outputs may create unwarranted confidence in the resulting analysis ^16–18^. As artificial intelligence makes increasingly sophisticated analyses accessible through natural-language interaction, the need for reliable domain-specific scientific software becomes more pressing rather than less. AI-assisted electrophysiological research therefore requires computational environments in which model-generated instructions are constrained by supported operations, methodological decisions remain subject to researcher oversight, and analytical execution can be independently traced ^19^.

A related challenge is the separation between methodological knowledge and computational practice. Analytical choices are often informed by previous publications, laboratory conventions or software defaults, yet the evidence underlying these choices is not necessarily preserved alongside their computational implementation ^19,20^. Even when methods are reported in detail, reproducing an analysis may require reconstructing software dependencies, parameter settings and processing decisions distributed across publications and software environments. Community initiatives such as EEG-BIDS and MEG-BIDS have improved data organization and methodological transparency, while reporting recommendations have established clearer expectations for reproducible electrophysiological research^21–23^. However, standardized data organization and methodological reporting do not, by themselves, ensure that an analytical workflow can be reconstructed and executed ^24,25^. Addressing this gap requires closer integration of published methodological evidence, computational configuration and persistent records of analytical execution. Importantly, agreement among published studies can inform methodological decisions without establishing that commonly reported procedures are necessarily optimal^26,27^.

Here we introduce MLE EEG Toolbox, an integrated framework for EEG and MEG analysis, AI-assisted workflow configuration, and computational reproducibility. Rather than developing new signal-processing algorithms, the framework integrates established methods into a unified environment where analytical decisions can be specified, reviewed, and preserved. It supports resting-state and task-based EEG analysis, MEG signal characterization, and source reconstruction. A locally deployed language-model assistant translates natural-language instructions into constrained, researcher-approved operations, while a literature-guidance module links analytical parameters to published methodological evidence. An integrated provenance system records processing decisions, software configurations, and execution histories to enable workflow reconstruction and replay. Implemented in interoperable MATLAB and standalone Java environments, the framework provides accessible analytical workflows without requiring extensive programming expertise. We evaluated its performance through cross-platform numerical comparisons and end-to-end analyses of public EEG and MEG datasets, demonstrating the integration of spectral, temporal, network, multivariate, and source-imaging methods. By combining AI-assisted interaction with controlled computation and systematic provenance tracking, MLE EEG Toolbox provides a practical foundation for accessible, transparent, and reproducible electrophysiological research. More broadly, it demonstrates how specialized scientific software can bridge flexible AI-generated instructions and reliable, traceable scientific computation.

## Results

### An integrated framework for electrophysiological analysis

MLE EEG Toolbox provides an integrated computational environment for electrophysiological research, connecting data preprocessing, signal characterization, statistical analysis, machine learning, and workflow documentation within a unified framework (Fig. 1). The platform supports commonly used electrophysiological data formats and organizes analytical procedures into configurable workflows, allowing researchers to move from raw recordings to processed data, quantitative features, statistical results, and visualizations without repeatedly switching between independent software environments.

**Figure 1.**
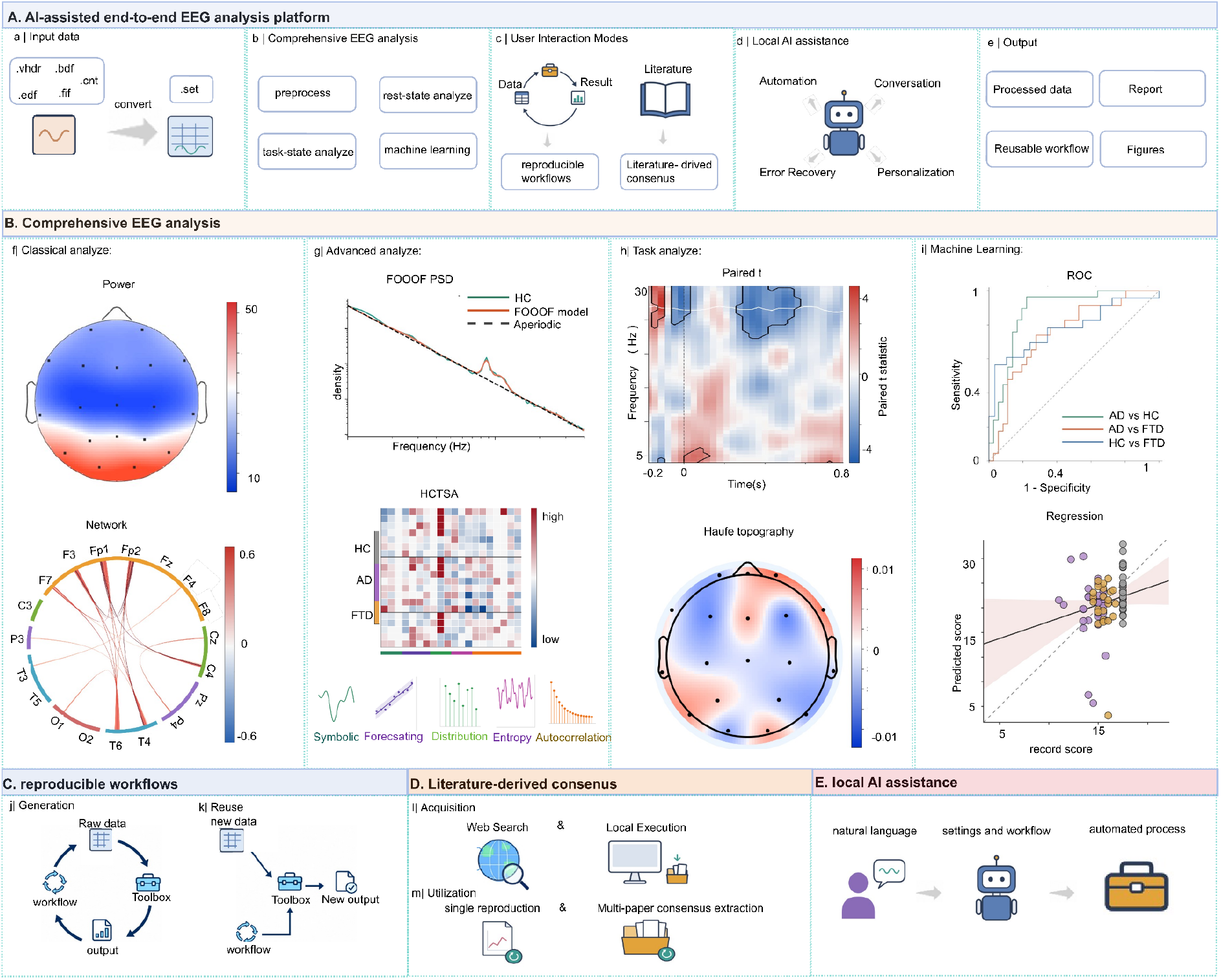
| Overview of the MLE EEG Toolbox and its integrated analytical framework. (A) AI-assisted end-to-end EEG analysis platform. (a) Data import and conversion from commonly used electrophysiological formats into a standardized dataset representation. (b) Integrated analytical modules covering preprocessing, resting-state analysis, task-related analysis, and machine learning. (c) User interaction modes supporting reproducible computational workflows and literature-informed parameter selection. (d) A locally deployed AI assistant supporting natural-language interaction, workflow automation, error recovery, and personalized analysis. (e) Standardized outputs, including processed datasets, analytical reports, figures, and reusable workflows. (B) Comprehensive EEG analysis. (f) Conventional EEG analysis, illustrated by scalp power topographies and functional connectivity networks. (g) Advanced analysis, including power spectral density decomposition using FOOOF and multivariate time-series characterization using highly comparative time-series analysis (HCTSA). (h) Task-related analysis, illustrated by event-related time-frequency representations and scalp topographies of statistical effects. (i) Machine-learning applications, including classification performance evaluated using receiver operating characteristic (ROC) curves and regression analysis comparing predicted and observed outcomes. (C) Reproducible workflows. (j) Automatic recording of processing operations, analytical parameters, and outputs during workflow generation. (k) Reuse of stored computational workflows to reproduce the analysis on new datasets. (D) Literature-derived consensus. (l) Acquisition of relevant scientific literature through online searches and local document processing. (m) Extraction and aggregation of methodological parameters to support single-study reproduction and cross-study consensus generation. (E) Local AI assistance. Natural-language instructions are translated into executable analysis plans through a locally deployed AI assistant, enabling user-supervised automation within the graphical interface.

The framework combines conventional electrophysiological methods with advanced analytical approaches. Classical analyses include spectral power estimation, scalp topography, functional connectivity, and network characterization, whereas advanced modules incorporate spectral parameterization, microstate analysis, and highly comparative time-series analysis (HCTSA). Task-related analyses support event-related responses, time–frequency decomposition, statistical comparisons, and multivariate decoding. The resulting features can be further integrated into supervised and unsupervised machine-learning workflows (Fig. 1A,B).

Beyond analytical functionality, the platform incorporates three complementary components designed to improve methodological accessibility and reproducibility: a locally deployed AI assistant for natural-language workflow configuration, a literature-guidance module for evidence-informed parameter selection, and a computational provenance system for recording and reproducing analytical operations (Fig. 1C–E). The MATLAB implementation integrates established electrophysiological analysis libraries, while the standalone Java application provides an alternative execution environment without requiring MATLAB. Both implementations employ compatible workflow representations and interoperable output formats.

The principal contribution of the framework is therefore not the introduction of individual signal-processing algorithms, but their integration into a shared configuration, execution, and provenance infrastructure that links analytical operations with their methodological decisions and computational histories.

### A constrained local AI assistant enables supervised workflow automation

MLE EEG Toolbox incorporates a locally deployed AI assistant that translates natural-language instructions into structured analytical operations, extracts methodological evidence from published literature, and uses analysis reports to outline a research framework (Fig. 2A). For workflow configuration, the assistant maps requests onto registered toolbox functions with predefined parameters, execution requirements, and validation rules. It uses the current graphical user interface (GUI) context to select relevant operations, configure their settings, and order them according to their dependencies.

**Figure 2.**
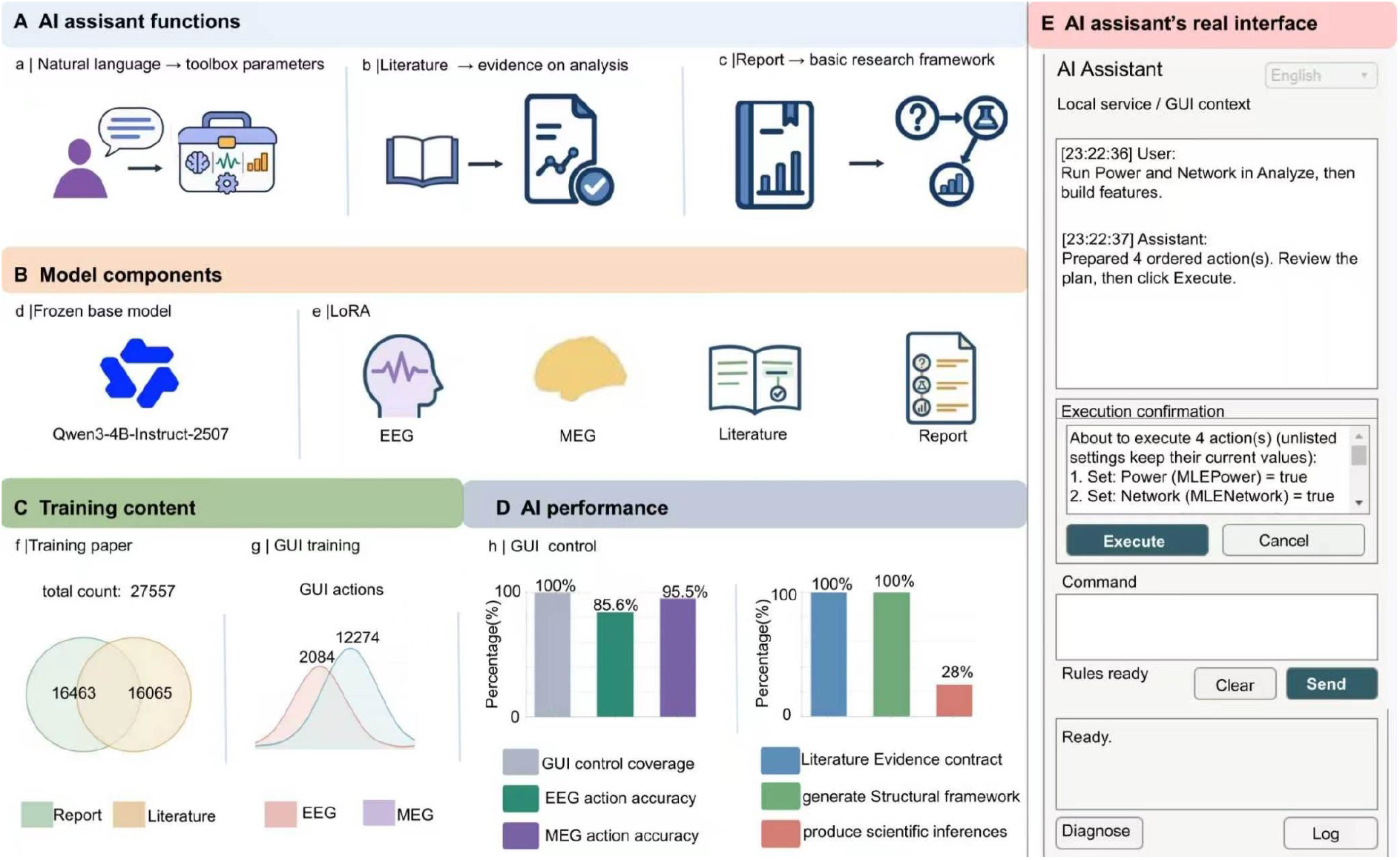
| Functions, model architecture, training data, and interface of the locally deployed AI assistant. (A) Assistant functions. The assistant translates natural-language instructions into toolbox parameters and graphical user interface (GUI) actions (a), extracts methodological evidence from the literature to inform analysis choices (b), and uses analysis reports to outline a research framework (c). **(B) Model architecture.** The assistant combines a frozen Qwen3-4B-Instruct-2507 base model (d) with task-specific low-rank adaptation (LoRA) modules for EEG and MEG workflow control, literature evidence extraction, and report-based research framing (e). Inference runs locally without per-request cloud-model token charges. **(C) Training data.** The literature and report modules used 27,557 unique papers, with 16,065 assigned to literature evidence extraction and 16,463 to the report task, including papers shared across tasks (f). GUI training used 2,084 EEG and 12,274 MEG action examples (g). **(D) Performance evaluation.** The assistant achieved 100% coverage of the evaluated GUI functions, with exact-action accuracies of 85.6% for EEG and 95.5% for MEG (h, left). In separate evaluations, the reported scores were 100% for literature evidence extraction, 100% for research framework generation, and 28% for scientific inference (h, right). **(E) Assistant interface.** In the example shown, a request to run power and network analyses and then build features is translated into a four-action plan. The interface displays the proposed operations and allows the researcher to review, approve, or cancel the plan before execution.

The assistant uses a frozen Qwen3-4B-Instruct-2507 base model with task-specific low-rank adaptation (LoRA) modules for EEG and MEG workflow control, literature evidence extraction, and report-based research framework generation (Fig. 2B). GUI training used 2,084 EEG and 12,274 MEG action examples. The literature and report tasks drew on 27,557 unique papers, with 16,065 used for literature evidence extraction and 16,463 for the report task, including papers shared between the two tasks (Fig. 2C).

In the GUI evaluation, the assistant covered 100% of the evaluated functions and achieved exact-action accuracies of 85.6% for EEG and 95.5% for MEG (Fig. 2D, left). The reported scores for literature evidence extraction, research framework generation, and scientific inference were 100%, 100%, and 28%, respectively (Fig. 2D, right). The 28% score highlights the need for researcher interpretation of analytical results.

In the example shown in Fig. 2E, a request to run power and network analyses and then build features is translated into a four-action plan. The toolbox checks the proposed parameters and dependencies and displays the ordered operations and their settings for researcher review. Execution requires explicit approval, and the researcher can cancel the plan before it runs. Inference runs locally, keeping electrophysiological recordings and natural-language instructions on the user’s machine and avoiding per-request cloud-model token charges.

### Literature-informed methodological guidance supports parameter selection and study reproduction

To connect analytical configuration with published methodological evidence, the literature-guidance module supports publication retrieval, parameter extraction, single-study protocol reconstruction, and cross-study consensus analysis (Fig. 3).

**Figure 3.**
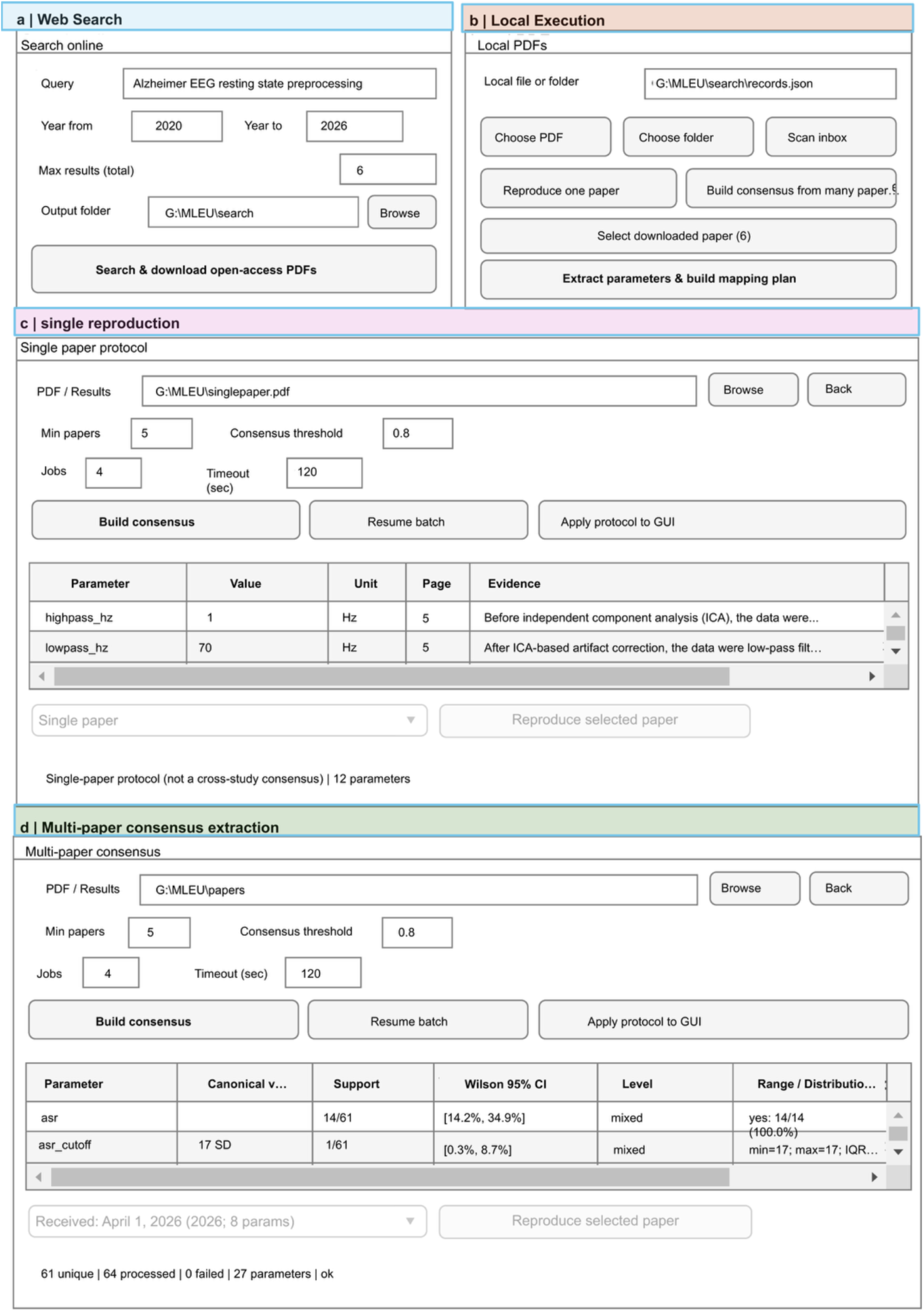
| Literature-guided methodological extraction, study reproduction, and cross-study consensus generation. (a) Web-based literature acquisition. Interface for searching and downloading open-access scientific publications using user-defined keywords, publication years, and retrieval limits. (b) Local literature processing. Interface for selecting locally stored PDF documents or literature collections, extracting methodological information, and generating structured parameter mappings for subsequent analysis. (c) Single-study methodological reproduction. Extraction of analysis parameters from an individual publication, with supporting evidence linked to the corresponding source-page locations. Extracted information is organized into a structured protocol that can be mapped to the graphical analysis interface to support reproduction of the reported analytical procedures. (d) Multi-study methodological consensus. Aggregation of extracted parameters across multiple publications to identify commonly reported analysis settings and characterize methodological variability. The interface displays canonical parameter values, study-level support, Wilson 95% confidence intervals for reporting proportions, and distributions of extracted settings. User-defined support thresholds facilitate the construction of literature-informed analysis protocols.

The literature acquisition interface enables researchers to search for relevant publications using user-defined keywords, publication periods, and retrieval limits. The system supports searches through PubMed, Europe PMC, OpenAlex, arXiv, and Crossref, with open-access full-text documents retrieved where permitted. Alternatively, researchers can provide locally stored PDF documents for direct methodological extraction (Fig. 3a,b).

For individual publications, the module extracts reported analytical parameters, including filter settings, sampling frequencies, referencing procedures, artifact-rejection criteria, independent component analysis configurations, and epoch definitions. Each extracted parameter is linked to its source document and supporting text, allowing researchers to inspect the evidence underlying the reconstructed analytical protocol. The resulting parameter mappings can subsequently be transferred to the toolbox interface to support reproduction of the reported processing procedures (Fig. 3c).

For collections of publications, the module aggregates extracted methodological information to characterize reporting frequencies and between-study variability (Fig. 3d). For each candidate parameter, the system summarizes the reported values, supporting publication counts, and distributions of methodological settings. Reporting proportions are accompanied by Wilson 95% confidence intervals to quantify uncertainty in the observed level of cross-study agreement. Configurable support thresholds allow researchers to distinguish frequently reported settings from parameters with limited or heterogeneous evidence.

The implementation also supports a more stringent classification of reporting consensus, requiring sufficient publication coverage, agreement exceeding a predefined threshold, and a lower Wilson confidence bound of at least 0.70. These criteria describe agreement within the retrieved literature rather than identifying universally optimal analytical settings. Accordingly, extracted recommendations remain subject to researcher review before they are incorporated into an executable workflow.

By linking proposed analysis parameters to their original published evidence, the module provides a traceable mechanism for literature-informed methodological configuration and reduces reliance on undocumented parameter selection.

### Computational provenance and workflow replay support reproducible electrophysiological analysis

MLE EEG Toolbox incorporates computational provenance directly into workflow execution, enabling analytical procedures to be recorded, inspected, and reproduced (Fig. 4). Rather than reconstructing processing histories retrospectively, the framework captures completed operations together with their resolved parameter values, input-file information, execution dependencies, and relevant software metadata.

**Figure 4.**
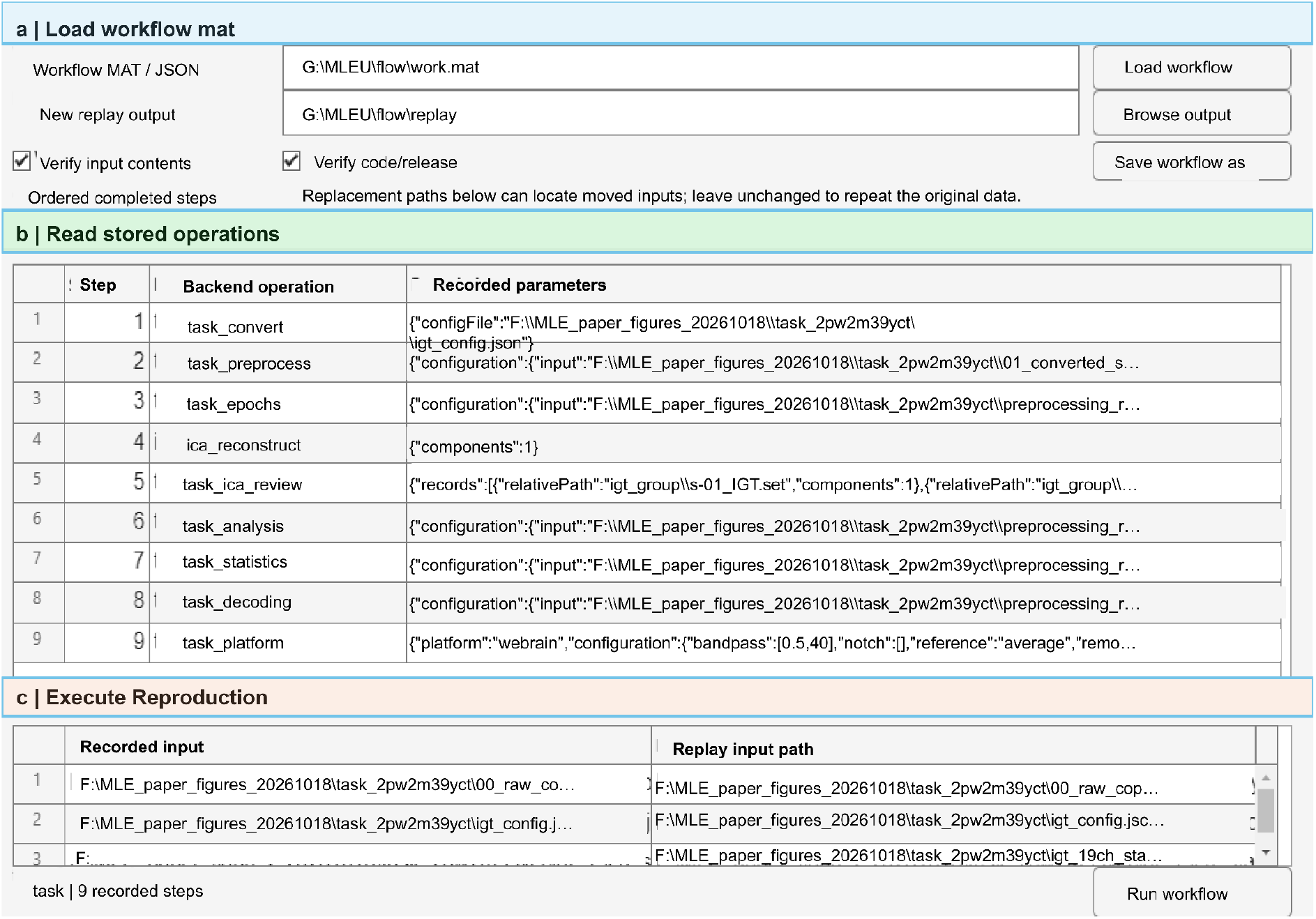
| Workflow provenance, parameter tracking, and reproducible execution. (a) Loading recorded computational workflows. Interface for importing previously saved workflow files in MAT or JSON format, specifying the destination for reproduced outputs, and enabling verification of input contents and software code or release information. (b) Inspection of recorded operations. Ordered processing history displaying completed computational steps, backend operation identifiers, and their associated parameter configurations. The illustrated task-analysis workflow includes data conversion, preprocessing, epoch extraction, independent component analysis (ICA) reconstruction and review, task analysis, statistical testing, decoding, and subsequent processing operations. (c) Workflow reproduction. Interface for mapping recorded input files to their corresponding locations in a new execution environment and replaying the stored computational operations. By preserving operation order and parameter configurations, the system supports reproducible execution, computational auditing, and transfer of analytical workflows across datasets and computing environments.

Recorded workflows can be saved in structured MAT or JSON formats and subsequently loaded into the workflow reproduction interface (Fig. 4a). The system presents the ordered computational history, including backend operation identifiers and the corresponding configurations. As illustrated in Fig. 4b, a representative task-processing workflow preserves successive operations covering data conversion, preprocessing, epoch extraction, ICA reconstruction and review, task analysis, statistical testing, and multivariate decoding.

To support execution in another computational environment, the replay interface allows researchers to inspect the original input references and assign replacement file locations when necessary (Fig. 4c). Input integrity can be checked using SHA-256 content fingerprints, while verification of software code or release information helps identify potential differences between the original and replay environments. Recorded parameters are recovered from the stored execution history rather than inferred from current GUI settings.

The system distinguishes two forms of repeated execution. In same-data replay, the original analytical configuration is reapplied to the corresponding input data, with available integrity checks used to verify file consistency. In protocol transfer, the recorded analysis configuration is applied to another dataset, producing a separate execution history and output directory. This distinction separates computational reproduction of an existing analysis from methodological reuse across independent datasets.

The provenance infrastructure therefore supports analytical traceability, workflow inspection, and controlled re-execution. Although these mechanisms improve computational reproducibility, they do not guarantee numerically identical results across different hardware architectures, software versions, or numerical implementations.

### Comprehensive resting-state EEG analysis integrates spectral, network, and temporal characterization

The resting-state analysis pathway integrates conventional electrophysiological measures with advanced characterization of neural signal dynamics and machine-learning applications (Fig. 5). To illustrate these capabilities, representative EEG analyses were performed across healthy control (HC), Alzheimer’s disease (AD), and frontotemporal dementia (FTD) groups.

**Figure 5.**
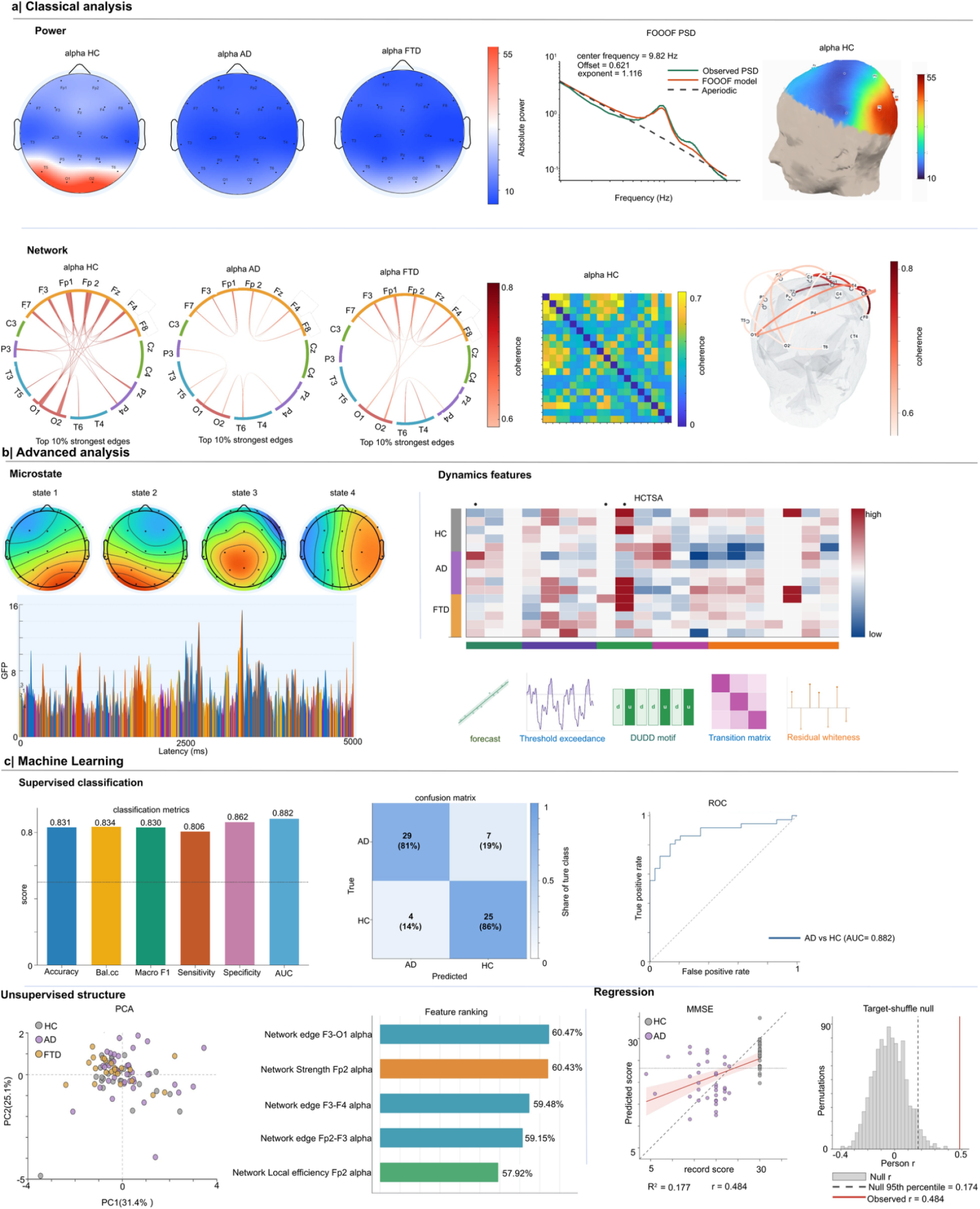
| Comprehensive resting-state EEG analysis and machine-learning capabilities. (a) Classical EEG analysis. Representative scalp topographies illustrating spatial distributions of alpha-band power across healthy controls (HC), Alzheimer’s disease (AD), and frontotemporal dementia (FTD) groups. Spectral characterization using FOOOF separates periodic oscillatory components from aperiodic background activity, with an example of estimated alpha peak characteristics. Cortical surface visualization illustrates the spatial distribution of EEG-derived spectral measures. Functional connectivity analysis is illustrated through thresholded network diagrams, connectivity matrices, and three-dimensional network representations. (b) Advanced EEG analysis. Representative microstate topographies and temporal sequences characterize recurring configurations of scalp electrical activity and their temporal organization. Highly comparative time-series analysis (HCTSA) provides a multidimensional characterization of signal dynamics, illustrated by standardized feature matrices across diagnostic groups and examples of temporal descriptors. (c) Machine-learning analysis. Supervised classification performance is summarized using accuracy, balanced accuracy, macro-F1 score, sensitivity, and area under the ROC curve (AUC). Confusion matrices and pairwise ROC curves illustrate discrimination among HC, AD, and FTD groups. Unsupervised analysis is demonstrated using principal component analysis (PCA) to visualize the distribution of participants in a reduced-dimensional feature space. Feature ranking identifies influential EEG-derived predictors, while regression analysis compares predicted and observed scores. A permutation-based null distribution provides a reference for evaluating the observed regression association.

Conventional spectral analyses generated scalp topographies of alpha-band power, illustrating differences in the spatial distribution of oscillatory activity across the example groups (Fig. 5a). Power spectral density estimates were further decomposed into periodic and aperiodic components using FOOOF, allowing oscillatory peak characteristics to be examined separately from the broadband spectral background. The framework also supports cortical visualization of EEG-derived measures and functional connectivity analysis using network diagrams, connectivity matrices, and three-dimensional representations.

Beyond conventional spectral and connectivity analysis, the advanced processing modules characterize the temporal organization of ongoing electrophysiological activity. Microstate analysis identifies recurring scalp potential configurations and their temporal expression, illustrated by representative spatial maps and state sequences (Fig. 5b). HCTSA provides complementary characterization through a broad collection of time-series features capturing properties such as temporal dependence, predictability, distributional structure, and signal variability. The resulting multidimensional feature profiles can be organized and visualized across participants or experimental groups.

Features derived from these analytical modules can subsequently enter machine-learning workflows (Fig. 5c). Supervised classification is illustrated through performance metrics, confusion matrices, and receiver operating characteristic (ROC) curves for discrimination among HC, AD, and FTD groups. Unsupervised dimensionality reduction using principal component analysis (PCA) provides a low-dimensional representation of participant-level feature variation, whereas feature-ranking procedures identify influential electrophysiological predictors. Regression functionality is demonstrated through comparisons between predicted and observed outcomes, together with a permutation-based reference distribution for evaluating predictive associations.

Collectively, these examples demonstrate the ability of the toolbox to combine established EEG measures, multidimensional temporal features, and predictive modelling within a common analytical environment. The illustrated group differences and predictive results are presented as functional demonstrations rather than independent evidence of diagnostic validity.

Task-related EEG analysis supports event-related responses, time–frequency decomposition, and multivariate inference

The task-related pathway provides an integrated workflow for characterizing stimulus-locked electrophysiological responses, condition-dependent spectral activity, statistical effects, and time-resolved decoding (Fig. 6).

**Figure 6.**
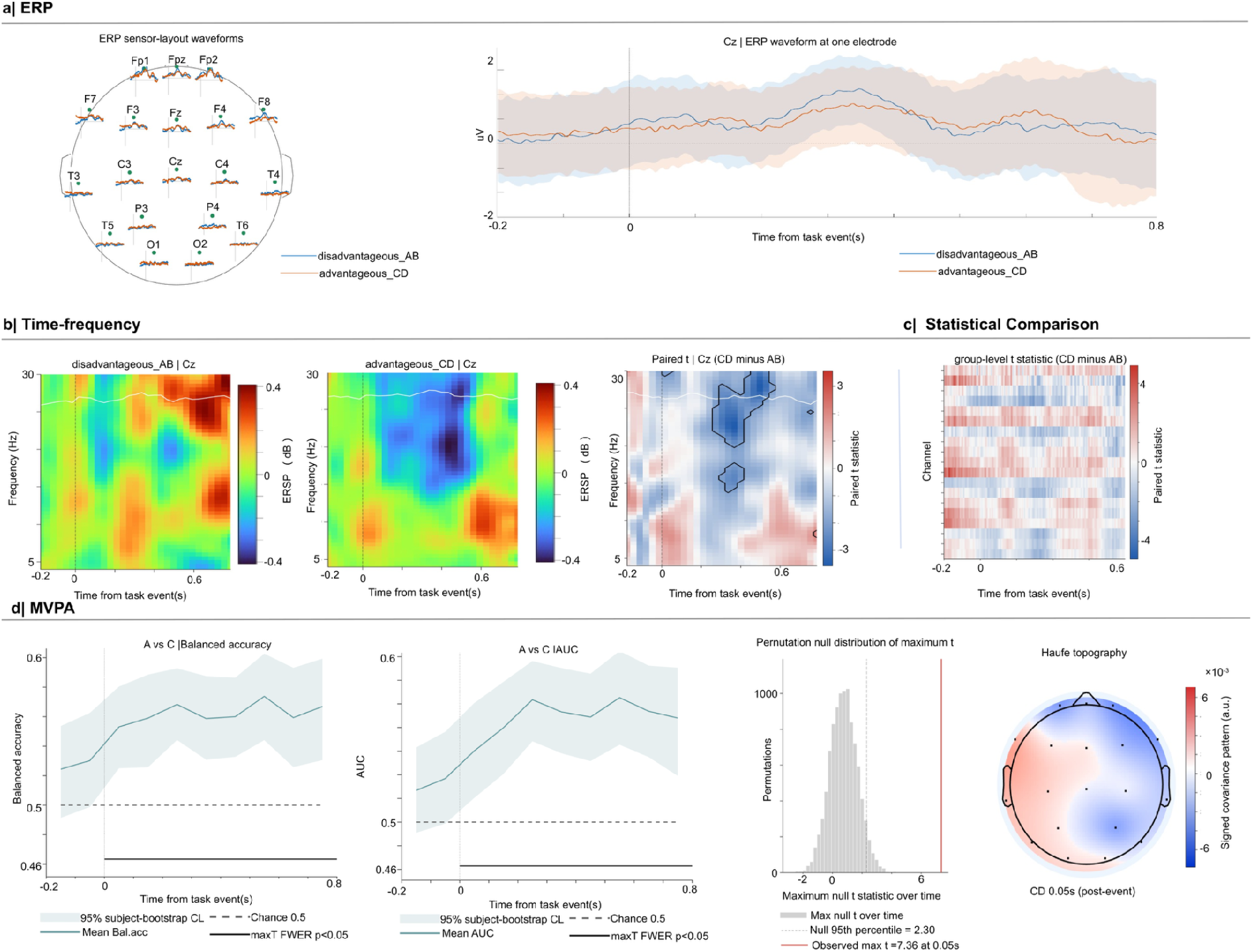
| Task-related EEG analysis, time-frequency decomposition, and multivariate statistical inference. (a) Event-related potential analysis. Representative event-related potential (ERP) waveforms across scalp electrodes and experimental conditions, together with an enlarged waveform visualization at a selected electrode. These analyses characterize the temporal evolution and spatial distribution of stimulus-locked electrophysiological responses. (b) Time-frequency analysis. Event-related spectral perturbation (ERSP) maps illustrating changes in oscillatory power across time and frequency for four experimental conditions (A–D). The representations provide a time-resolved characterization of task-related spectral modulations. (c) Statistical comparison. Time-frequency statistical map showing condition-related differences using paired-sample t statistics. Contours identify time-frequency regions meeting the specified statistical significance criterion. (d) Multivariate pattern analysis (MVPA). Time-resolved decoding performance assessed using area under the ROC curve (AUC), with shaded regions representing 95% subject-bootstrap confidence intervals. A permutation-based null distribution of the maximum t statistic across time points is used for family-wise error control, with the observed maximum statistic compared against the 95th percentile of the null distribution. A scalp topography illustrates the spatial distribution of decoding-related effects.

Following event identification and epoch extraction, the framework computes event-related potentials (ERPs) across electrodes and experimental conditions. Representative waveforms illustrate the temporal evolution of evoked responses across the scalp, with selected electrodes displayed in greater detail to facilitate comparison among experimental conditions (Fig. 6a).

To characterize frequency-specific responses, the toolbox applies time–frequency decomposition and generates event-related spectral perturbation (ERSP) maps. The representative analysis displays changes in spectral power across time and frequency for four experimental conditions, illustrating the capacity to examine both transient and sustained oscillatory responses during task performance (Fig. 6b).

Condition-dependent electrophysiological differences can be assessed using statistical comparisons across the time–frequency domain. The example paired-sample analysis produces a map of *t* statistics, with contours highlighting regions satisfying the specified statistical criterion (Fig. 6c). This provides a spatially and temporally resolved representation of task-related spectral differences.

The framework further supports multivariate pattern analysis (MVPA) to quantify the information carried by electrophysiological signals over time. In the illustrated decoding analysis, classifier performance is represented as a time-resolved area under the ROC curve (AUC), with 95% subject-bootstrap confidence intervals (Fig. 6d). Statistical inference is supported by a permutation-based distribution of the maximum *t* statistic across time, providing a reference for family-wise error control. Decoding-related scalp topographies additionally illustrate the spatial distribution of the corresponding electrophysiological effects.

These analyses demonstrate the integration of univariate signal characterization, time–frequency statistics, and multivariate decoding within a continuous task-processing workflow.

### Integrated MEG analysis extends the framework to source-resolved neural dynamics

To demonstrate applicability beyond EEG, the framework also supports MEG analysis, integrating sensor-level responses, spectral characterization, functional connectivity, machine-learning classification, and source reconstruction (Fig. 7).

**Figure 7.**
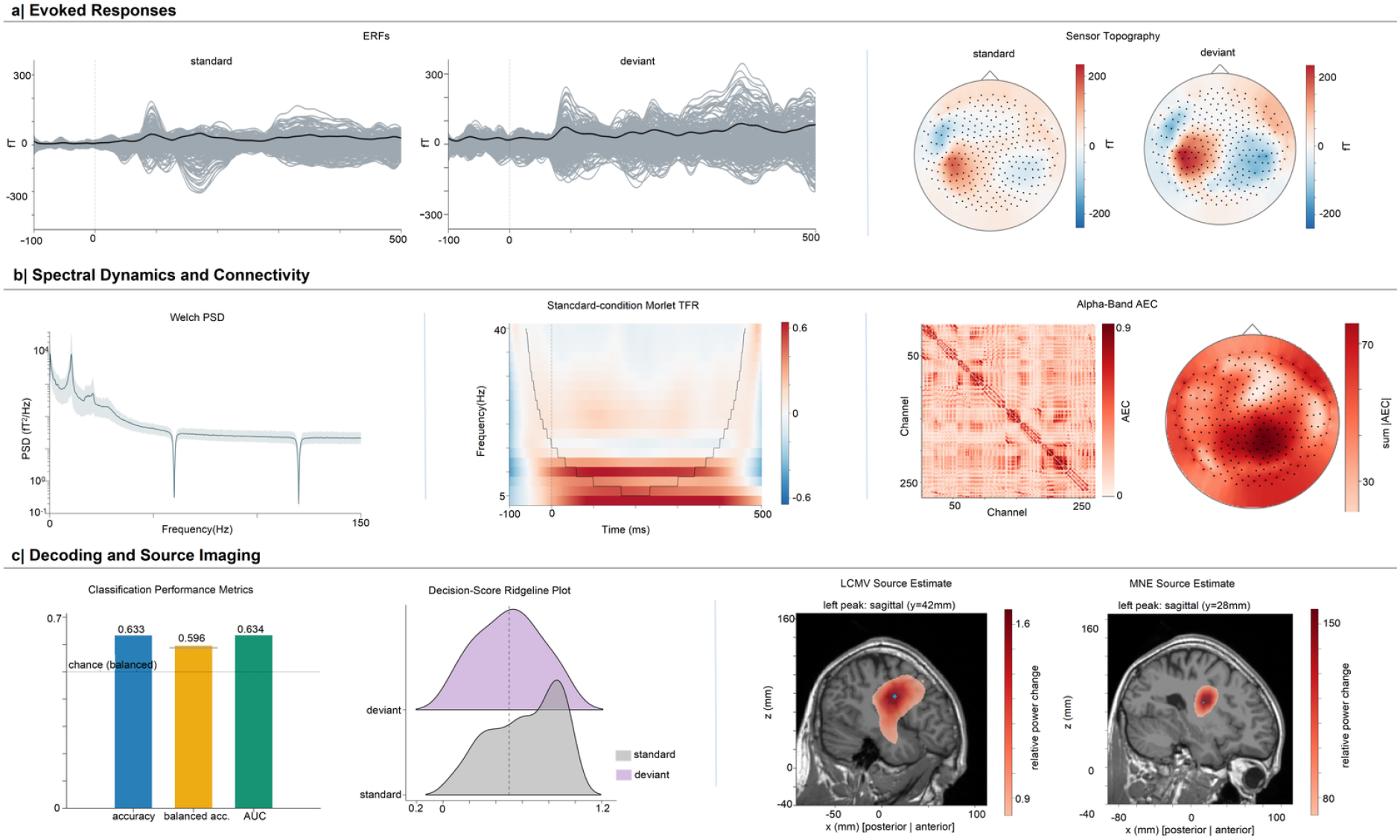
| Comprehensive MEG analysis integrating evoked responses, spectral dynamics, functional connectivity, machine learning, and source imaging. (a) Evoked responses. Event-related magnetic fields (ERFs) elicited by standard and deviant stimuli, showing sensor-level temporal responses across MEG channels (grey traces) and their averaged responses (black traces). Sensor topographies illustrate the spatial distributions of evoked magnetic fields for the two experimental conditions. (b) Spectral dynamics and functional connectivity. Power spectral density (PSD) estimated using Welch’s method, illustrating the frequency-dependent organization of spontaneous MEG activity. Morlet wavelet-based time-frequency representation (TFR) showing stimulus-related spectral power changes in the standard condition. Alpha-band amplitude envelope correlation (AEC) analysis, illustrated by a sensor-level functional connectivity matrix and a corresponding scalp topography summarizing connectivity strength across sensors. (c) Decoding and source imaging. Machine-learning classification of standard and deviant conditions, with performance summarized using accuracy, balanced accuracy, and area under the receiver operating characteristic curve (AUC). Ridge-density plots illustrate the distributions of classifier decision scores for the two conditions. Source reconstruction results obtained using linearly constrained minimum variance (LCMV) beamforming and minimum-norm estimation (MNE) are displayed on representative sagittal anatomical slices, illustrating the spatial distribution of estimated source activity. Colour scales indicate relative power changes for the two reconstruction methods.

Event-related magnetic fields (ERFs) were examined for standard and deviant stimulus conditions. Sensor-level time courses revealed stimulus-locked magnetic responses with distinct temporal profiles across the two conditions, while topographic representations illustrated their spatial distributions across MEG sensors (Fig. 7a). These analyses demonstrate the ability to characterize both temporal and spatial aspects of event-related magnetic activity.

Spectral analysis using Welch’s method generated power spectral density estimates, capturing the frequency-dependent structure of ongoing MEG signals. Morlet wavelet-based time–frequency analysis further illustrated stimulus-related spectral dynamics in the standard condition (Fig. 7b). Functional connectivity was characterized using alpha-band amplitude envelope correlation (AEC), yielding a sensor-by-sensor connectivity matrix and a corresponding topographic representation of aggregate connectivity strength. Together, these analyses demonstrate complementary characterization of spectral and network properties at the sensor level.

The framework additionally supports multivariate decoding of experimental conditions. In the representative standard-versus-deviant classification, accuracy, balanced accuracy, and AUC reached 0.633, 0.596, and 0.634, respectively (Fig. 7c). Distributions of classifier decision scores further illustrated differences in the model outputs between the two conditions. These results provide an example of condition-related information captured by the implemented decoding pipeline, without establishing generalizable classification performance beyond the evaluated data.

Finally, anatomically informed source reconstruction was demonstrated using linearly constrained minimum variance (LCMV) beamforming and minimum-norm estimation (MNE). The resulting source estimates were visualized on sagittal anatomical images, illustrating the spatial distribution of reconstructed neural activity and the distinct patterns produced by the two inverse approaches (Fig. 7c). These examples demonstrate that the framework can integrate sensor-level analysis and source-space estimation within the same computational environment.

Together, the MEG demonstrations extend the analytical scope of the platform from EEG-based characterization to source-resolved electrophysiological analysis, while retaining the same general framework for computational execution and workflow documentation.

### Cross-platform validation and public-data analyses establish computational feasibility

To evaluate computational consistency between software implementations, we conducted 142 automated cross-platform tests comparing the Java and MATLAB versions under matched input data and analytical configurations. Core spectral and connectivity operations showed close numerical agreement, with relative errors below (10^{-10}) for spectral power estimates and a maximum absolute difference of (4.2 \times 10^{-15}) for phase-locking values. The evaluated Brain Connectivity Toolbox graph metrics produced identical outputs, while six machine-learning classifiers generated identical predictions and scores across implementations.

Small implementation-dependent differences were observed for selected analytical procedures. FOOOF peak-frequency estimates differed by up to 0.36 Hz in 6 of 32 evaluated channels. Differences in microstate coverage did not exceed 0.004, while mean microstate durations differed by no more than 0.11 ms. Because ICA decomposition can be sensitive to initialization, convergence criteria, and numerical implementation, equivalent but not necessarily numerically identical decompositions were expected across platforms.

End-to-end feasibility was further evaluated using publicly available EEG recordings. Resting-state processing was examined using OpenNeuro dataset ds004504, comprising 88 participants. Task-based analysis was evaluated using an Iowa Gambling Task dataset containing 55 participants and 7,590 valid trials. The task-processing workflow supported event identification, epoch-based analysis, feature generation, and multivariate decoding. Linear discriminant analysis with within-participant cross-validation achieved an AUC of 0.5438, with a permutation-test *P* value of 0.0050.

The cross-platform comparisons establish numerical consistency for the evaluated operations and configurations, while the public-data applications demonstrate that the framework can execute complete analytical workflows on independent electrophysiological datasets. These results support the computational feasibility of the integrated platform but do not independently establish the scientific validity of AI-generated plans, literature-derived recommendations, or the generalizability of the illustrated machine-learning models.

## Discussion

The increasing sophistication of electrophysiological analysis has created a growing gap between the availability of computational methods and the ability to apply them transparently and reproducibly. Advances in artificial intelligence may narrow this gap by allowing researchers to interact with scientific software through natural language, but they also introduce new challenges concerning methodological validity, computational accountability and researcher oversight. Here, we developed the MLE Toolbox to address these interconnected challenges through an integrated framework combining established electrophysiological methods, constrained AI-assisted workflow configuration, literature-informed parameter selection and persistent computational provenance. Across EEG and MEG applications, we demonstrate the integration of conventional and advanced analytical procedures, while cross-platform comparisons establish numerical consistency for a range of core operations. Together, these results provide a practical example of how AI-assisted scientific analysis can be embedded within a structured computational environment in which methodological decisions and analytical execution remain explicit and traceable.

An important implication of this work is that the emergence of AI-assisted research increases, rather than diminishes, the need for reliable domain-specific scientific software^27,28^. Large language models can interpret research objectives and generate plausible analytical procedures, but linguistic competence does not establish the scientific appropriateness of the resulting analyses^28^. Electrophysiological research is particularly sensitive to this distinction because preprocessing decisions, signal representations, statistical assumptions and experimental designs are tightly interdependent^25,26^. An analytically plausible sequence of operations may nevertheless alter the signals of interest, introduce data leakage or produce misleading statistical inferences^28–30^. MLE EEG Toolbox addresses part of this challenge by separating natural-language interpretation from computational execution. The AI assistant operates through a constrained catalogue of registered functions, while parameter validation, dependency checks and researcher approval provide additional boundaries on its actions. This design does not eliminate methodological errors, but it establishes a clearer separation between model-generated recommendations and the scientific computations used to implement them^31,32^. More broadly, it suggests that the value of AI in computational neuroscience may depend less on its ability to generate unrestricted analytical procedures than on how effectively those procedures can be constrained, inspected and evaluated within reliable scientific environments.

The performance of the local AI assistant provides an initial assessment of this approach. In the reported evaluation, the assistant covered all evaluated GUI functions and achieved exact-action performance of 85.6% for EEG operations and 95.5% for MEG operations. These findings indicate that natural-language requests can be mapped to predefined analytical operations with substantial, although incomplete, accuracy. Importantly, successful execution of an interface action is not equivalent to selecting a scientifically appropriate analysis^32^. The distinction is especially consequential when requests are ambiguous, when several valid analytical approaches are available or when the experimental design imposes constraints that are not fully specified in the user instruction^33,34^. Human supervision therefore remains an essential component rather than a temporary limitation of the framework. The assistant is intended to reduce the technical burden of configuring and coordinating analyses, not to replace the scientific judgement required to select methods, interpret results or evaluate their validity^35^. Future benchmarks should move beyond action-level accuracy to assess whether AI-generated plans preserve experimental constraints, avoid inappropriate analytical choices and enable researchers to identify and correct errors before execution.

A second contribution is the integration of published methodological evidence into the configuration of computational workflows. EEG analysis frequently depends on choices that are informed by previous studies but remain only loosely connected to the resulting computational implementation^27,35^. By extracting reported parameters from the literature, preserving their source evidence and summarizing reporting frequencies across publications, MLE EEG Toolbox provides a mechanism for making the evidential basis of these decisions more explicit. The distinction between reproducing an individual published protocol and identifying common practices across studies is particularly important^34^. Single-study reconstruction aims to recover the methodological choices of a specific investigation, whereas cross-study aggregation characterizes how frequently particular procedures are reported within a body of literature. Neither approach establishes that the extracted parameters are optimal for a new experiment. Widely adopted settings may reflect historical conventions, default software configurations or characteristics of particular recording systems. Moreover, literature retrieval and automated extraction can introduce biases that are not captured by confidence intervals around reporting proportions^35^. Accordingly, the literature-guidance module is best understood as a tool for methodological transparency and evidence-informed decision-making, rather than an automated authority for parameter selection. Independent evaluation against expert-annotated publications will be necessary to determine its extraction accuracy and practical reliability.

Computational provenance provides a complementary foundation for reproducible research by preserving how analytical decisions are implemented rather than relying exclusively on retrospective methodological descriptions. Conventional reporting standards and structured data formats have improved the documentation and exchange of electrophysiological recordings, but the ability to identify inputs and describe processing steps does not necessarily guarantee that an analysis can be reconstructed and executed^25^. In MLE EEG Toolbox, computational operations are recorded together with their resolved parameters, dependencies, input-data fingerprints and relevant software information. This approach connects methodological configuration to the actual execution history and allows individual processing steps to be inspected or replayed. The distinction between same-data replay and protocol transfer further separates two objectives that are often grouped under the term reproducibility: reconstructing a computation using its original inputs and applying an established analytical procedure to an independent dataset. Both are important for cumulative neuroscience, but they address different sources of uncertainty. Moreover, provenance records cannot fully eliminate numerical differences arising from software versions, hardware architectures, parallel computation or stochastic optimization. Their principal value is therefore to make computational differences identifiable and investigable rather than to guarantee identical outputs under all conditions.

Several limitations qualify the interpretation of the present findings. First, the evaluation primarily establishes software functionality, numerical consistency and the feasibility of integrated electrophysiological analysis. Although the framework is designed to improve accessibility and reproducibility, the present results do not demonstrate that it reduces analysis time, prevents methodological errors or improves scientific outcomes relative to existing software environments. Controlled studies involving researchers with different levels of programming and electrophysiological expertise will be required to evaluate these potential benefits. Second, the AI assistant has been evaluated primarily at the level of supported operations and action selection. More demanding assessments should examine its responses to ambiguous or incomplete instructions, its rejection of unsupported analyses, its adherence to experimental constraints and the effectiveness of researcher oversight. Third, the literature-guidance module requires independent validation of parameter extraction, numerical value matching and source attribution, particularly for publications with incomplete reporting or heterogeneous methodological terminology. Fourth, computational provenance is limited to operations represented within the framework and cannot automatically recover undocumented processing decisions made before data import. Finally, although the platform supports diverse EEG and MEG analyses, the present demonstrations do not establish clinical diagnostic validity, and their scientific interpretation remains dependent on appropriate experimental design, statistical evaluation and independent replication.

Future developments should therefore prioritize validation of the scientific reliability of AI-assisted workflows rather than simply expanding the range of automated functions. Particularly informative evaluations would compare constrained and unconstrained AI-generated analysis plans, measure the frequency and consequences of methodological errors, and determine whether provenance-aware workflows improve the ability of independent researchers to reconstruct published analyses. Benchmarking literature-derived parameter recommendations against expert assessments would also clarify when automated methodological guidance is reliable and when additional human interpretation is required. Extending these evaluations across software environments, electrophysiological modalities and independent datasets would help determine the generalizability of the framework. Such assessments would move beyond demonstrating that complex analyses can be automated towards establishing whether AI-assisted scientific workflows can be made more dependable in routine research practice.

In conclusion, MLE EEG Toolbox provides an integrated framework linking methodological knowledge, researcher oversight and computational execution in electrophysiological research. Its central contribution is not a new measure of brain activity, but a computational architecture in which established analytical methods can be configured through constrained AI assistance, informed by published evidence and documented through persistent execution records. As artificial intelligence becomes more closely integrated into scientific research, the challenge is not merely to make sophisticated analyses easier to initiate, but to ensure that they remain methodologically justified, computationally traceable and open to independent scrutiny. The present framework represents a step towards this goal and illustrates a broader role for domain-specific scientific software as an essential foundation for accountable AI-assisted neuroscience.

## Methods

### Software architecture and implementation

MLE EEG Toolbox was developed as an integrated computational framework for electrophysiological data analysis, combining graphical workflow configuration, established signal-processing methods, artificial intelligence (AI)-assisted analysis planning, literature-informed methodological guidance, and computational provenance tracking. The framework comprises two interoperable implementations: a MATLAB-based research platform and a standalone Java desktop application. Both implementations support project configuration, data preprocessing, analytical execution, visualization, and workflow management, while providing complementary computational environments for electrophysiological research.

The MATLAB implementation supports MATLAB R2020b and later releases and integrates established EEG analysis libraries within a modular scientific computing environment. The Java application operates independently of MATLAB and incorporates a locally deployed language-model assistant, literature-guidance functions, and workflow execution infrastructure. Interoperability is supported through a shared workflow representation and structured provenance records, together with commonly used electrophysiological and computational data formats, including EEGLAB SET/FDT, MAT, JSON, and CSV.

The software architecture separates workflow configuration, computational execution, and provenance management into interconnected components. Researchers define analytical procedures through graphical controls or AI-assisted instructions, which are translated into registered operations and executed by the corresponding computational backend. Each operation is associated with an explicit parameter configuration and execution history. Compatibility of workflow representations is distinguished from numerical equivalence of computational implementations; agreement between corresponding MATLAB and Java operations was therefore assessed empirically rather than assumed.

### Electrophysiological data import and project configuration

The toolbox supports commonly used electrophysiological data formats, including EEGLAB SET, BrainVision, EDF/BDF, Neuroscan CNT, FIF, and MAT^22,23^. During project initialization, recording files are associated with participant identifiers, experimental conditions or diagnostic groups, and available sensor metadata. These associations establish a structured relationship between electrophysiological recordings and the analytical procedures subsequently applied to them.

For external input files, SHA-256 cryptographic hashes are generated and retained within the computational provenance records. These fingerprints provide a means of verifying file identity and detecting changes to the original inputs during subsequent workflow execution or replay. Data import and project configuration are recorded as explicit processing steps to maintain traceability between source recordings, analysis inputs, and generated outputs.

### EEG preprocessing

EEG preprocessing is organized into three configurable pathways: manually specified processing, automated artifact cleaning, and protocol-defined processing^8,36^. These pathways provide different levels of automation while retaining access to the underlying analytical parameters^36–40^.

Available preprocessing operations include DC offset removal, digital filtering, rereferencing, channel-quality assessment, bad-channel interpolation, and independent component analysis (ICA)^8,38,41,42^. Where supported by the computational backend, established procedures such as clean_rawdata and ICLabel are integrated to assist with artifact identification and component assessment. ICA components can be inspected through their spatial distributions, spectral characteristics, and temporal activity before researchers determine whether individual components should be retained or rejected. Preprocessing operations are executed using the parameters selected for the corresponding workflow. The system records the effective parameter values applied during execution, rather than reconstructing the configuration retrospectively from the graphical interface. This distinction allows preprocessing histories to be inspected and, where supported, reproduced using the stored analytical settings.

### Resting-state EEG analysis

The resting-state analysis module integrates conventional electrophysiological measures with complementary approaches for characterizing spectral, network, and temporal properties of neural activity.

Spectral analysis includes power spectral density estimation using Welch’s method. Functional connectivity can be quantified using phase-locking value (PLV), phase-lag index (PLI), and weighted phase-lag index (wPLI), with the resulting connectivity matrices available for network characterization^43–46^. Supported graph-theoretical measures include node strength and clustering coefficient.

The framework additionally supports spectral parameterization using FOOOF/specparam, which models the power spectrum in terms of periodic oscillatory components and an aperiodic background^47^. This representation allows oscillatory peak characteristics to be examined separately from broadband spectral structure. Temporal organization can be examined using EEG microstate segmentation based on modified k-means clustering^48^. The resulting microstate configurations and temporal sequences provide complementary descriptions of recurring scalp potential patterns. More extensive time-series characterization is supported through catch22 and, optionally, highly comparative time-series analysis (hctsa), enabling the extraction of features describing multiple statistical and dynamical properties of electrophysiological signals^49–51^.

These modules implement established analytical approaches. Their integration within MLE EEG Toolbox is intended to facilitate coordinated analysis and computational documentation rather than introduce new signal-processing estimators.

### Task-based EEG analysis

The task-based analysis pathway supports the processing of event-related electrophysiological recordings, from event identification and epoch extraction to statistical inference and multivariate decoding. Recorded event codes are identified and mapped to experimental conditions. Researchers configure epoch windows and baseline intervals before subsequent analysis. The workflow retains information concerning valid trials, rejected epochs, and participant-level quality-control outcomes to facilitate inspection of the processing history^8,23,52^. Event-related potentials (ERPs) are obtained from condition-specific epochs and can be visualized across electrodes and experimental conditions. Time-frequency decomposition is supported for examining changes in spectral activity during task performance, together with inter-trial phase coherence for characterizing the consistency of stimulus-related phase responses. Statistical analysis functions support comparisons of electrophysiological measures across experimental conditions. Multivariate pattern analysis enables time-resolved decoding of condition-related information from electrophysiological features^25,29,41,53^. The specific statistical procedures and validation schemes are selected according to the experimental design and inferential objective, with the corresponding analytical settings retained in the workflow record.

### Machine-learning analysis

The machine-learning module supports supervised and unsupervised analysis of features derived from resting-state or task-based electrophysiological recordings^54,55^.

Available classification methods include linear discriminant analysis (LDA), support vector machines (SVMs), logistic regression, naive Bayes, k-nearest neighbours, decision trees, random forests, ensemble methods, and neural networks. The framework also provides regression methods for continuous outcome prediction and dimensionality-reduction approaches for exploratory characterization of high-dimensional feature spaces.

Model evaluation can incorporate cross-validation, classification performance metrics, and permutation testing. Supported performance measures include accuracy, balanced accuracy, sensitivity, F1 score, and area under the receiver operating characteristic curve (AUC), as appropriate for the prediction task. Regression performance can be evaluated through comparisons between observed and predicted outcomes.

Cross-validation procedures are configured according to the intended generalization target. For analyses seeking prediction across independent participants, training and test partitions are separated at the participant level to avoid information leakage between observations belonging to the same individual. Data-dependent preprocessing, feature selection, and model tuning should be restricted to the training data within the corresponding validation procedure to preserve the independence of held-out observations.

Permutation testing provides an empirical null distribution against which observed predictive performance can be evaluated. The interpretation of predictive results depends on the sampling design, validation scheme, and independence of the evaluation data.

### Locally deployed AI assistant and model architecture

MLE EEG Toolbox incorporates a locally deployed AI assistant based on Qwen3-4B-Instruct-2507, with a frozen pretrained backbone and task-specific low-rank adaptation (LoRA) modules for EEG and MEG workflow control, literature evidence extraction, and report-based research framework generation^56,57^.The assistant was trained using natural-language instructions paired with registered GUI operations and scientific literature to support analysis configuration, methodological guidance, and research planning (Supplementary Methods).

The assistant translates natural-language requests and current GUI context into structured analytical operations defined within the toolbox. Each proposed operation is subject to parameter and dependency validation and requires researcher approval before execution. The literature-related modules extract methodological information from publications and organize research reports to support evidence-informed analysis and workflow design. All inference runs locally within the standalone Java application without external AI APIs or per-request token charges. AI-assisted operations use the same execution and provenance infrastructure as manually configured workflows, enabling transparent, researcher-controlled analysis.

### Evaluation of AI-assisted operation selection

The AI assistant was evaluated for its ability to map natural-language instructions to registered GUI operations. The assessment included functional coverage and exact-action performance for EEG and MEG operations. Functional coverage was defined as the proportion of evaluated toolbox functions for which the assistant supported the corresponding operation. Exact-action performance quantified the proportion of evaluated instructions for which the generated action matched the expected registered operation. Performance was summarized separately for EEG and MEG operations. The evaluation assessed the assistant’s ability to interact with the constrained operational interface and did not independently establish the scientific validity of complete analysis plans or their suitability for particular experimental designs.

### Literature retrieval and methodological parameter extraction

The literature-guidance module was designed to connect analytical configuration with methodological information reported in published studies.

The module retrieves bibliographic records from five sources: PubMed, Europe PMC, OpenAlex, arXiv, and Crossref. Search parameters can be specified by researchers, and retrieved records are deduplicated using publication identifiers and bibliographic metadata. Openly accessible full-text PDF documents are obtained where available and permitted. The system also supports the analysis of locally provided PDF documents. Methodological extraction focuses on parameters commonly used in electrophysiological research, including filtering frequencies, sampling rates, referencing schemes, artifact-rejection criteria, ICA methods, epoch definitions, and baseline intervals. Extracted information is organized into structured parameter records containing the reported value, associated units, supporting source-text excerpt, and document location. These records allow researchers to inspect the provenance of extracted parameters and assess their correspondence with the original publication. The module supports two complementary applications. In single-study protocol reconstruction, methodological parameters are extracted from an individual publication and organized into a candidate analysis configuration. In cross-study aggregation, parameter records from multiple publications are combined to characterize reported methodological practices and identify settings supported by the retrieved literature. Extracted parameters are not applied automatically solely on the basis of their reporting frequency. Researchers can inspect, accept, modify, or reject candidate settings before they are incorporated into executable workflows.

### Quantification of methodological reporting consistency

To characterize agreement in methodological reporting across studies, the literature-guidance module calculates the proportion of eligible publications supporting a specified parameter value or range. For a candidate parameter reported by k of n eligible publications, the observed reporting proportion was calculated as p = k/n. Uncertainty in the observed reporting proportion was estimated using the Wilson score 95% confidence interval, with the standard normal critical value set to z = 1.96. The interval was calculated from the observed proportion and the number of eligible publications. Under the current configuration, a parameter was classified as exhibiting strong reporting consensus when at least 30 eligible publications contributed evidence, the observed support proportion exceeded a configurable threshold, and the lower bound of the Wilson confidence interval was at least 0.70. Parameters with insufficient publication coverage or weaker evidence were classified separately. These estimates quantify consistency of methodological reporting within the retrieved literature. They do not measure the accuracy of automated parameter extraction, account for potential selection biases in literature retrieval, or establish that a frequently reported parameter is scientifically optimal. The resulting summaries were therefore presented as evidence-informed guidance rather than definitive methodological recommendations.

### Computational provenance and workflow replay

The framework incorporates computational provenance directly into analytical execution to preserve the relationship between methodological configuration, processing operations, and generated outputs. Each completed operation generates a structured record containing its operation identifier, resolved parameter values, input-file fingerprints, upstream dependencies, relevant software or function versions, and, where applicable, random-number generator states. Analytical outputs and visualizations are linked to the operations responsible for their generation. Workflow records are stored in structured formats, including MAT and JSON, and can be loaded into a dedicated replay interface. The interface provides an ordered representation of previously executed operations and their parameter configurations. Successive executions are recorded in separate output directories to preserve earlier analytical results and facilitate comparison between runs.

Two complementary replay modes are supported. Same-data replay aims to reproduce an analysis using the original recordings and associated inputs. Before execution, available file fingerprints are checked against the stored records to identify changes in the input data. Protocol transfer preserves a recorded analytical configuration while allowing researchers to supply a different dataset. Input compatibility and subsequent execution outcomes are recorded separately from the original workflow. This mode supports the reuse of established analysis procedures across independent datasets without treating the resulting outputs as repetitions of the original computation. The provenance infrastructure is designed to support computational auditing and reproducibility assessment. Recording analytical parameters, input identities, and software information does not guarantee numerical identity across different computing environments, particularly for procedures involving stochastic initialization, numerical optimization, or implementation-dependent floating-point operations.

### Cross-platform numerical validation

Numerical consistency between the MATLAB and Java implementations was evaluated through 142 automated cross-platform tests using matched input data and analytical configurations. The evaluation included operations covering digital filtering, spectral estimation, functional connectivity, graph-theoretical characterization, and machine learning. For corresponding operations implemented in both environments, output agreement was evaluated using metrics appropriate to the resulting data structure.

For numerical outputs, absolute differences between corresponding Java and MATLAB estimates were calculated. Relative differences were used where appropriate to assess the magnitude of numerical discrepancies relative to the reference output. For classification procedures, agreement was evaluated through comparisons of predicted labels and associated prediction scores. Methods involving optimization or stochastic initialization were considered separately because equivalent analytical configurations do not necessarily yield identical numerical solutions. Implementation-dependent deviations were retained and reported rather than treated as evidence of complete equivalence. This evaluation was designed to assess computational consistency between corresponding implementations under the tested conditions. It was not intended to establish the clinical validity or neurophysiological specificity of the implemented analytical measures.

### Validation using public EEG and MEG datasets

End-to-end workflow execution was evaluated using three publicly available electrophysiological datasets representing resting-state EEG, task-based EEG, and MEG. These datasets were selected to examine the applicability of the framework across recording modalities, experimental designs, and analytical procedures.

### Resting-state EEG dataset

Resting-state EEG analyses were demonstrated using the publicly available dataset *A Dataset of EEG Recordings from Alzheimer’s Disease, Frontotemporal Dementia and Healthy Subjects* (OpenNeuro accession ds004504). The dataset comprises eyes-closed resting-state recordings from 88 participants, including 36 individuals with Alzheimer’s disease (AD), 23 with frontotemporal dementia (FTD), and 29 healthy controls (HC). The dataset was used to demonstrate the execution of spectral analysis, functional connectivity estimation, network characterization, microstate analysis, time-series feature extraction, and machine-learning procedures. Representative outputs included power topographies, spectral parameterization results, connectivity matrices, microstate representations, and multivariate feature-based analyses. The purpose of this application was to evaluate the ability of the framework to coordinate heterogeneous analytical procedures within an integrated workflow. Group differences and classification examples were not intended to establish new diagnostic biomarkers or independently validate the clinical utility of the implemented measures.

### Task-based EEG dataset

Task-based EEG processing was evaluated using the publicly available *Iowa Gambling Task (IGT) and EEG Data* dataset, comprising 55 participants and 7,590 valid trials. The dataset was used to demonstrate event-related processing, including event identification, epoch extraction, ERP estimation, time-frequency analysis, statistical comparisons, and multivariate decoding. For the reported single-trial classification example, linear discriminant analysis was performed using within-participant cross-validation. Predictive performance was evaluated using the area under the receiver operating characteristic curve, and statistical significance was assessed using permutation testing. The decoding analysis was treated as a demonstration of computational workflow execution rather than evidence of a newly identified task-related neural signature. Its interpretation was restricted to the evaluated dataset, prediction procedure, and validation scheme.

### MEG dataset

MEG functionality was assessed using the publicly available Brainstorm MEG sample dataset (OpenNeuro accession ds000246). The dataset includes two auditory oddball recordings from a single participant acquired using a 275-channel whole-head MEG system. The analysis included sensor-level event-related magnetic fields (ERFs), power spectral density estimation using Welch’s method, Morlet wavelet-based time-frequency decomposition, and functional connectivity estimation using alpha-band amplitude envelope correlation (AEC). To demonstrate multivariate analysis, standard and deviant stimulus conditions were evaluated using a classification workflow, with performance summarized using accuracy, balanced accuracy, and AUC. Source reconstruction was demonstrated using linearly constrained minimum variance (LCMV) beamforming and minimum-norm estimation (MNE). Reconstructed activity was visualized on anatomical images to illustrate the integration of sensor-level measurements with source-space analysis.

Because the MEG example involved recordings from a single participant, it was used exclusively to evaluate functionality and workflow compatibility. The resulting sensor-level differences, decoding performance, and reconstructed source patterns were not interpreted as population-level neurophysiological findings. Together, the three datasets provided complementary test cases for evaluating the execution of integrated EEG and MEG workflows. These analyses demonstrate the computational scope and feasibility of the framework under the evaluated conditions without establishing the scientific validity of every supported analytical procedure.

## Acknowledgments

The authors acknowledge the developers of Brainstorm, FieldTrip, EEGLAB, MNE-Python, the Brain Connectivity Toolbox, and FreeSurfer, upon whose work MLE-Toolbox builds. We thank [funding agencies and colleagues] for support and feedback during toolbox development.

## Data and Code Availability

MLE-Toolbox source code, example datasets, and documentation are available at https://neuro-tool-works.pages.dev/. The toolbox is distributed under the GNU General Public License v3.0. Example datasets are provided under the Creative Commons Attribution 4.0 license.

## Supplementary Methods

### Locally deployed AI assistant and model architecture

MLE EEG Toolbox incorporates a locally deployed artificial intelligence (AI) assistant designed to translate natural-language research instructions into structured electrophysiological analysis operations, retrieve methodological evidence from scientific literature, and assist in organizing research workflows. The assistant is built on Qwen3-4B-Instruct-2507, a pretrained language model adapted to domain-specific tasks using low-rank adaptation (LoRA). The pretrained model parameters remain frozen, while task-specific adaptation components support EEG and MEG workflow control, literature evidence extraction, and report-based research framework generation (Fig. 2A,B).

The architecture separates natural-language interpretation from computational execution. Rather than allowing the language model to generate unrestricted executable code, the assistant maps researcher instructions onto a predefined catalogue of analytical operations registered within the toolbox. Each registered operation is associated with a function identifier, configurable parameters, admissible parameter values, and execution requirements. The language model uses the researcher’s instruction together with the current graphical user interface (GUI) context to identify relevant operations and propose their configurations. These proposals are subsequently interpreted and validated by the toolbox before being submitted to the corresponding computational backend.

The assistant also supports literature-informed methodological guidance. Task-specific language-model components extract analytical information from scientific publications and organize information from analysis reports into candidate research frameworks. These functions are intended to assist methodological interpretation and research planning rather than replace independent scientific evaluation.

Model inference is performed locally within the standalone Java application, without requiring access to external language-model APIs or per-request cloud-model token usage. Electrophysiological recordings, local file paths, and natural-language research instructions can therefore remain within the user’s computational environment. The assistant operates within the same workflow configuration and provenance infrastructure used for manually specified analyses, ensuring that AI-assisted and conventional operations remain subject to common validation and execution procedures.

### Construction of natural-language and GUI-operation training datasets

To adapt the language model to electrophysiological workflow control, task-specific training datasets were assembled to associate natural-language research instructions with analytical operations supported by MLE EEG Toolbox. The GUI-control training data comprised 2,084 EEG and 12,274 MEG action examples (Fig. 2C). These datasets were designed to represent the relationship between researcher instructions, the available GUI functions, and the analytical operations implemented by the corresponding computational backend.

The supported operations were defined through the toolbox’s registered function catalogue. Each operation specifies its analytical purpose, associated interface action, available parameters, and execution prerequisites. This representation provides a constrained set of possible targets for language-model-assisted workflow configuration, allowing the assistant to identify supported functions rather than construct arbitrary programmatic commands.

Natural-language instructions and relevant interface context served as inputs for action prediction. The corresponding targets represented the registered analytical operations required to satisfy the instructions. The EEG operation set covered commonly used procedures for data preprocessing, spectral characterization, functional connectivity, network analysis, time-series feature extraction, task-related analysis, and machine learning. The MEG operation set included sensor-level electrophysiological analysis, spectral and time–frequency characterization, functional connectivity, multivariate analysis, and source reconstruction.

The training framework was intended to support the interpretation of analytical requests expressed in natural language and their translation into operations compatible with the toolbox. During inference, operation selection remains constrained by the registered function definitions, while parameter validity and dependencies are checked independently of the language model. This separation is important because selecting an appropriate interface action does not by itself establish that its parameters or position within a processing workflow are scientifically valid.

### Preparation of literature-based training data

The literature-related components were developed using a corpus of 27,557 unique scientific publications. Of these, 16,065 publications were assigned to literature evidence extraction and 16,463 to report-based research framework generation, with overlap between the two task collections (Fig. 2C).

The literature evidence-extraction task focused on recovering methodological information relevant to electrophysiological research. Target information included filtering settings, sampling frequencies, referencing procedures, artifact-rejection criteria, independent component analysis configurations, epoch definitions, and other parameters reported in analytical methods. Extracted information was organized into structured methodological records that could be linked to supporting source passages and used to inform subsequent analysis configuration.

The report-based task supported the organization of information from research reports into structured descriptions of analytical objectives, methodological procedures, and potential research workflows. This functionality was designed to assist the formulation and organization of research analyses rather than independently determine the scientific validity of a proposed study.

These task-specific datasets supported the adaptation of the language model to methodological text and research-oriented instructions. During routine toolbox use, literature retrieval and document processing are performed through the literature-guidance module. Retrieved publications provide methodological evidence for inspection and parameter selection; their retrieval does not itself imply modification of the pretrained model or its task-specific adaptation parameters.

### Task-specific language-model adaptation

The Qwen3-4B-Instruct-2507 model was adapted to the electrophysiological research environment using task-specific LoRA components while retaining the pretrained backbone parameters. This approach enables task-specific adaptation without requiring full-parameter fine-tuning of the underlying language model.

The adaptation framework encompasses GUI operation control for EEG and MEG, literature evidence extraction, and report-based research framework generation. Each task defines a correspondence between the relevant input information and the expected output representation. For GUI control, the objective is to map natural-language requests and interface context to registered operations. For literature extraction, the objective is to recover structured methodological information from scientific text. For report-based assistance, the objective is to organize supplied research information into an interpretable analytical framework.

The use of a frozen backbone with task-specific adaptation separates the general language capabilities of the pretrained model from the specialized requirements of electrophysiological workflow configuration and methodological information processing. The resulting model components are integrated into the local assistant, allowing task-dependent interpretation of user instructions and research documents within a shared software environment.

### Constrained workflow generation and researcher-supervised execution

The AI assistant generates candidate analytical operations within the set of functions registered in MLE EEG Toolbox. When a researcher submits a natural-language instruction, the system interprets the request together with the current GUI context and identifies the analytical functions needed to address it. For instructions requiring multiple processing steps, the assistant proposes an ordered sequence of operations according to the analytical dependencies represented within the toolbox.

Each proposed action is associated with its corresponding function identifier and parameter configuration. Before execution, the toolbox checks whether the requested function is supported, whether the proposed parameter values satisfy the registered constraints, and whether the required input data and preceding operations are available. These checks are implemented within the computational framework rather than delegated entirely to the language model.

The resulting workflow is presented to the researcher for inspection. Researchers can review the proposed operations and their parameters, modify the configuration where supported, or cancel the plan before execution. Computational operations are initiated only after explicit researcher approval. This human-supervised design preserves control over methodological decisions while reducing the need to configure every supported analysis through separate interface interactions.

For example, a request to perform spectral power analysis, estimate functional connectivity, and subsequently construct electrophysiological features can be represented as a sequence of registered analytical operations (Fig. 2E). The system identifies the required functions, organizes their execution order, checks the associated parameters and dependencies, and presents the resulting plan for review.

The assistant also supports the identification of errors encountered during workflow execution and the proposal of recovery operations within the supported function catalogue. Recovery actions remain subject to the same operational constraints and, where required, researcher confirmation.

After execution, completed operations are recorded by the computational provenance infrastructure, including their resolved parameter values, input information, execution dependencies, and relevant software metadata. This allows AI-assisted workflows to be inspected and, where supported, replayed using the same provenance mechanisms as manually configured analyses.

#### Evaluation of AI-assisted operation selection and research-support tasks

The AI assistant was evaluated for its ability to translate natural-language instructions into registered EEG and MEG GUI operations, with performance assessed separately for the two modalities (Fig. 2D).

Functional coverage was defined as the proportion of evaluated toolbox functions for which the assistant supported the corresponding operation. Exact-action accuracy was defined as the proportion of evaluated instructions for which the predicted registered operation matched the expected operation. The assistant covered 100% of the evaluated GUI functions and achieved exact-action accuracies of 85.6% for EEG and 95.5% for MEG operations.

The literature-related components were additionally evaluated for methodological evidence extraction, research framework generation, and scientific inference. The reported evaluation scores were 100%, 100%, and 28%, respectively (Fig. 2D). These results were treated as task-specific evaluation outcomes rather than interchangeable measures of scientific reasoning or methodological validity. In particular, the substantially lower scientific inference score indicates that the ability to extract information or generate a structured research framework does not establish the reliability of independent scientific interpretation.

The GUI evaluation assessed the compatibility of model-generated actions with the toolbox’s constrained operational interface. Correct selection of a registered operation does not necessarily imply that the selected analysis is appropriate for a particular experimental design, that all parameter settings are scientifically justified, or that a complete multistep workflow is free from methodological errors. Similarly, successful methodological extraction or report organization does not independently establish the accuracy of the underlying scientific conclusions.

Accordingly, the AI assistant was designed as a researcher-supervised interface to established computational methods rather than an autonomous scientific decision-making system. Its evaluation characterizes the performance of the implemented assistance functions, while the scientific suitability of analytical choices remains subject to researcher assessment and independent validation.

**Supplementary Fig. 1.**
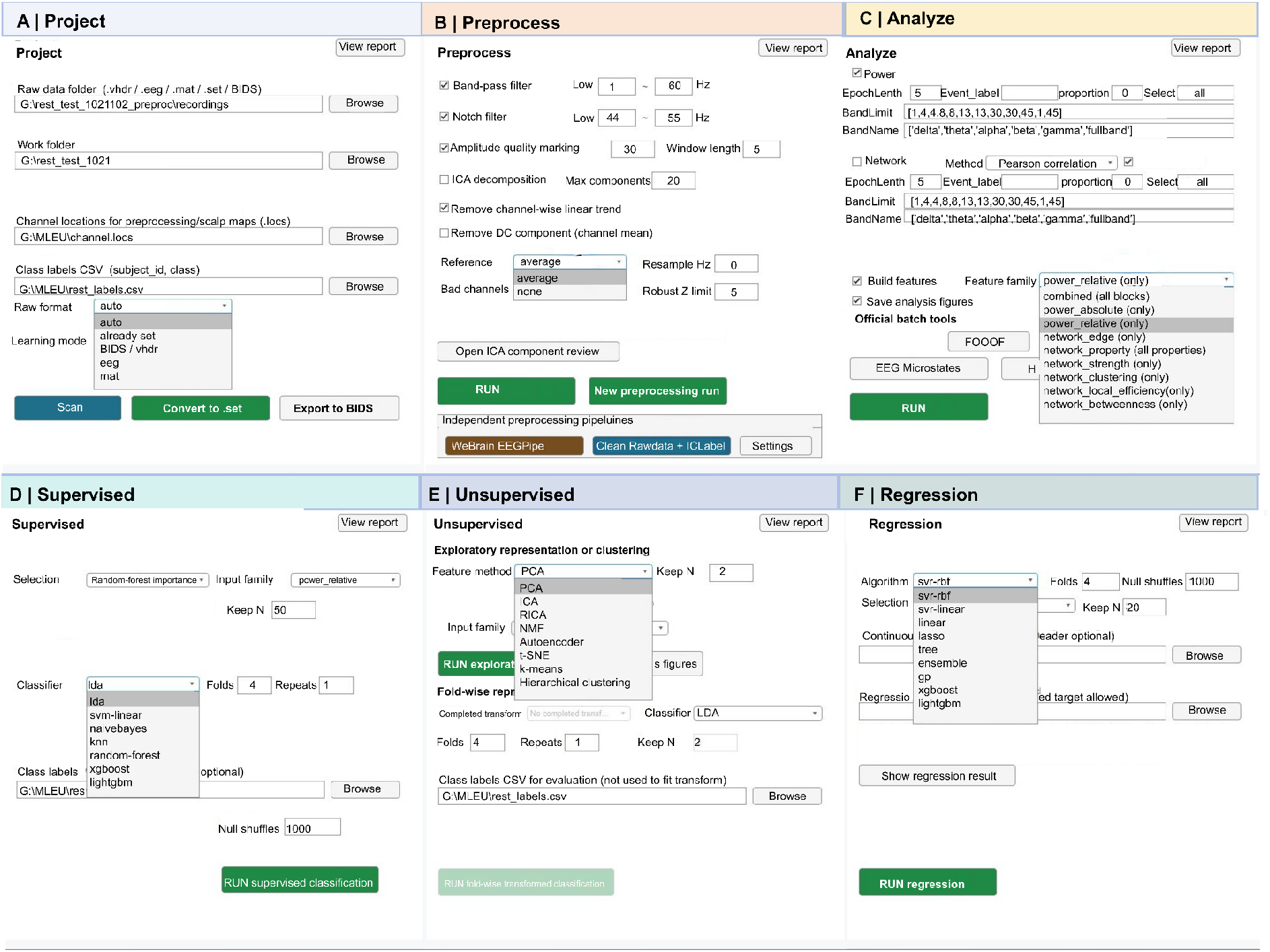
| Graphical user interface for resting-state EEG analysis and machine learning in MLE Toolbox. a, Project configuration interface for importing EEG recordings in multiple file formats, specifying working directories, electrode locations and participant-level class labels, and converting datasets to EEGLAB or BIDS formats. b, Preprocessing interface supporting band-pass and notch filtering, amplitude-based quality assessment, independent component analysis (ICA), detrending, re-referencing, resampling and bad-channel detection, together with alternative preprocessing pipelines. c, Analysis interface for spectral power estimation, functional connectivity, graph-theoretical characterization and feature construction, with integrated tools for spectral parameterization and EEG microstate analysis. d, Supervised learning interface for feature selection and classification using multiple machine-learning algorithms, with configurable cross-validation and permutation testing. e, Unsupervised learning interface for dimensionality reduction, latent representation learning and clustering, with support for fold-wise feature transformation and subsequent classification. f, Regression interface for predicting continuous outcomes from EEG-derived features using multiple regression algorithms, with configurable feature selection, cross-validation and permutation testing

**Supplementary Fig. 2.**
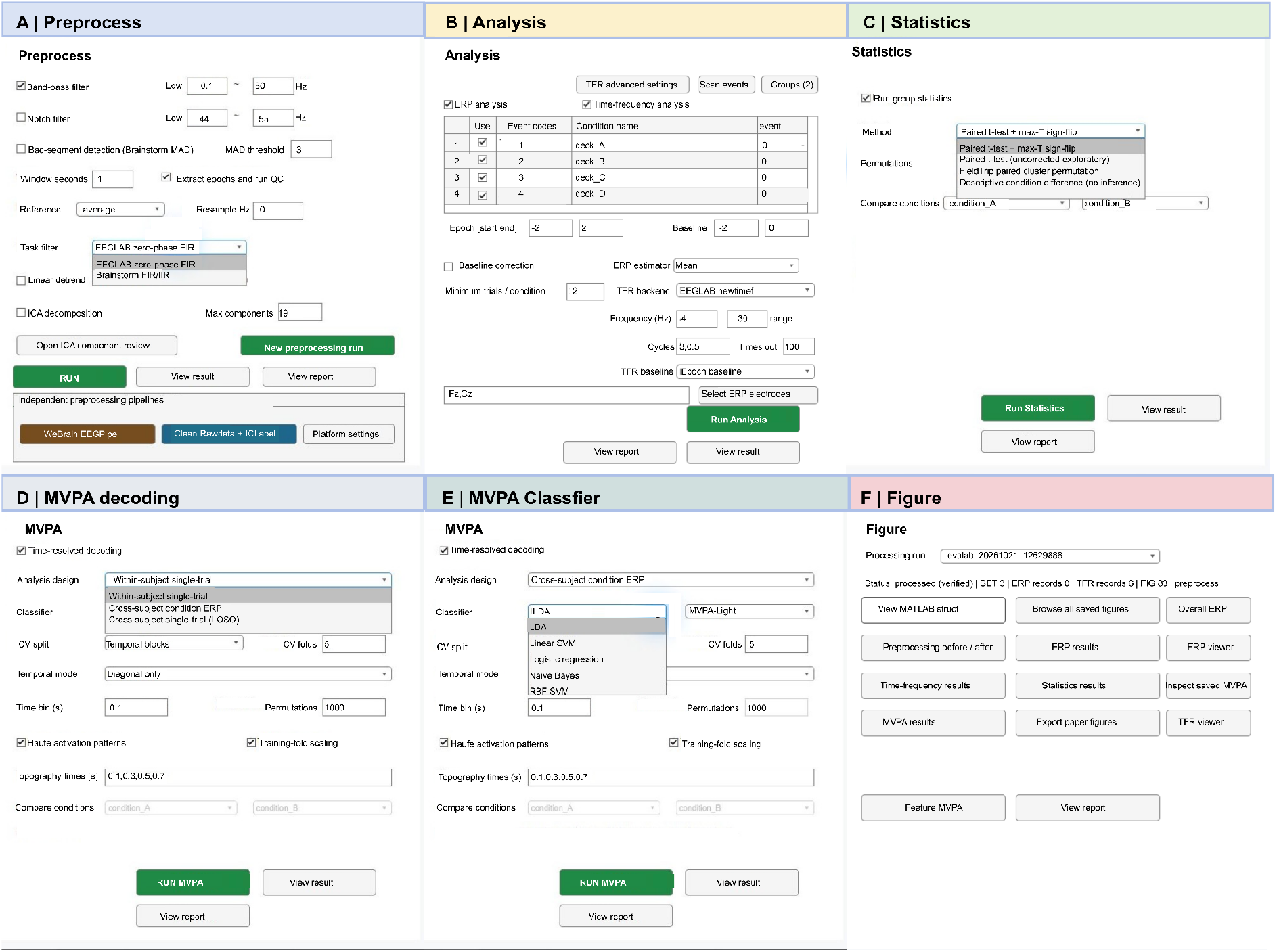
| Graphical user interface for task-based EEG analysis and multivariate decoding in MLE Toolbox. a, Preprocessing interface for task-based EEG recordings, including temporal filtering, bad-segment detection, epoch extraction, quality control, re-referencing, resampling and independent component analysis (ICA). b, Event-related analysis interface for defining experimental conditions, epoch boundaries, baseline intervals and parameters for event-related potential (ERP) and time–frequency analyses. c, Statistical analysis interface supporting paired comparisons, maximum-statistic permutation testing, cluster-based permutation testing and descriptive condition differences. d, Multivariate pattern analysis (MVPA) interface supporting within-subject single-trial decoding, cross-subject condition-level ERP decoding and cross-subject single-trial decoding with leave-one-subject-out validation. Configurable parameters include cross-validation strategies, temporal analysis modes, time-bin duration, permutation testing and Haufe-transformed activation patterns. e, Classifier configuration for time-resolved decoding, supporting linear discriminant analysis, linear support vector machines, logistic regression, naïve Bayes and radial basis function support vector machines, with integration of the MVPA-Light backend. f, Results visualization and management interface for inspecting preprocessing outcomes, ERP responses, time–frequency representations, statistical results and multivariate decoding performance, together with options for exporting publication figures and accessing processing reports.

**Supplementary Fig. 3.**
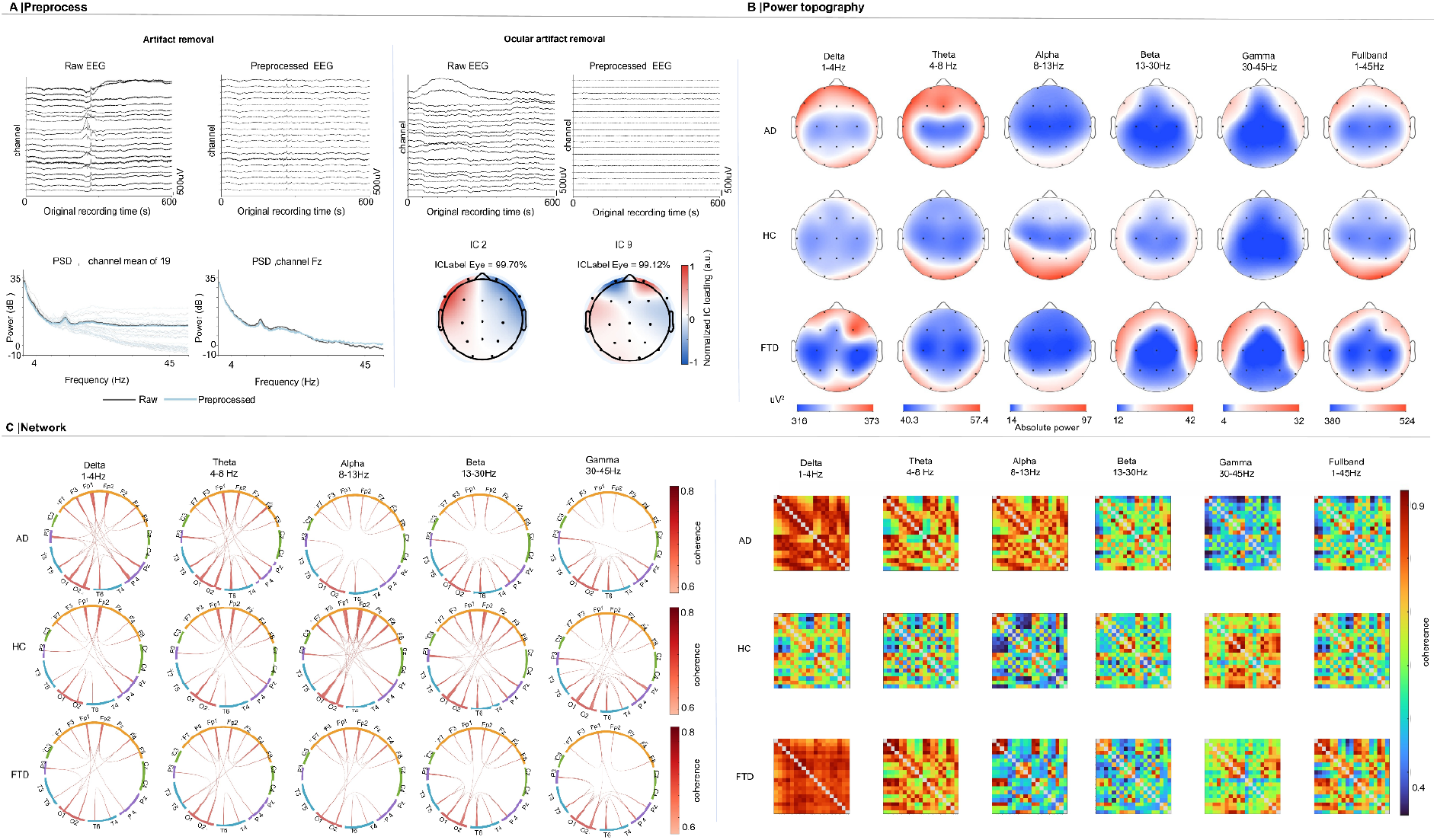
| Integrated preprocessing, spectral analysis and functional connectivity characterization of resting-state EEG. a, EEG preprocessing. Representative multichannel EEG recordings before (left) and after (right) preprocessing, illustrating the attenuation of prominent artifacts while preserving ongoing electrophysiological activity. The lower panels compare power spectral density (PSD) estimates before and after preprocessing, averaged across 19 EEG channels (left) and at the Fz electrode (right). b, Spectral power topography. Scalp distributions of EEG spectral power in individuals with Alzheimer’s disease (AD), healthy controls (HC) and frontotemporal dementia (FTD), displayed across delta (1–4 Hz), theta (4–8 Hz), alpha (8–13 Hz), beta (13–30 Hz), gamma (30–45 Hz) and broadband (1–45 Hz) frequency ranges. Colour scales represent spectral power (µV²), with separate ranges for each frequency band. c, Frequency-specific functional connectivity. Circular connectivity diagrams (left) and corresponding connectivity matrices (right) illustrating functional interactions between EEG channels in the AD, HC and FTD groups. Connectivity was characterized using the phase-locking value (PLV) across the same frequency ranges. Circular diagrams summarize the spatial organization and strength of interchannel connections, whereas connectivity matrices provide complementary representations of pairwise phase synchronization. These analyses demonstrate the integration of EEG preprocessing, spectral characterization and network-level analysis within MLE EEG Toolbox.

**Supplementary Fig. 4.**
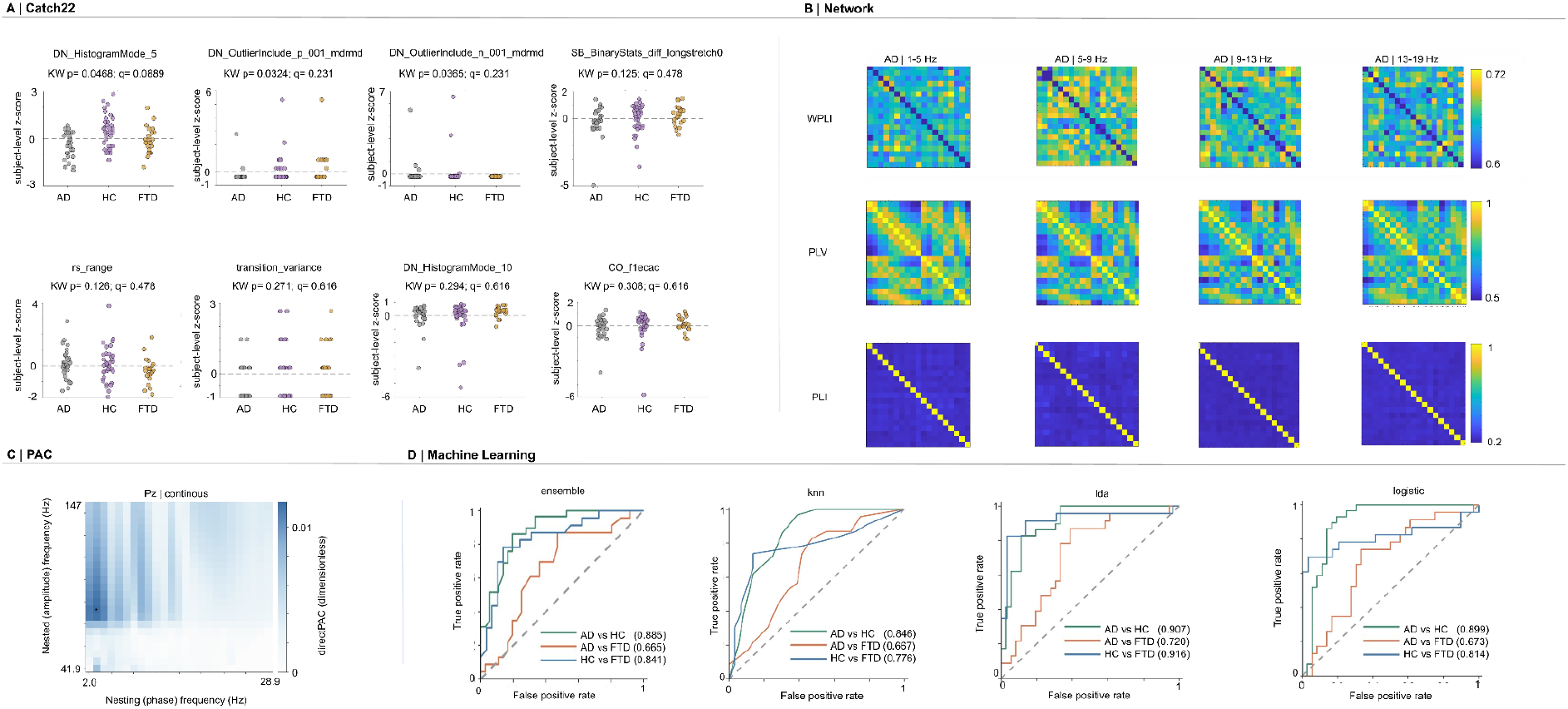
| Multidimensional EEG time-series analysis, cross-frequency coupling and machine-learning classification. **a, Time-series feature extraction.** Representative EEG time-series features derived from highly comparative time-series characterization, including measures of temporal variability, distributional properties, nonlinear complexity and long-range temporal dependence. Scatter plots show individual feature values for participants with Alzheimer’s disease (AD), healthy controls (HC) and frontotemporal dementia (FTD). Group differences were assessed using Kruskal–Wallis tests, with nominal P values and false discovery rate (FDR)-adjusted q values indicated above each feature. **b, Functional connectivity analysis using HERMES.** Representative weighted phase-lag index (wPLI) connectivity matrices obtained using the HERMES framework across four frequency bands (1–5, 5–9, 9–13 and 13–19 Hz) for the AD, HC and FTD groups. Matrix entries represent pairwise phase-based functional connectivity between EEG channels, with colour indicating wPLI magnitude. **c, Phase–amplitude coupling.** Representative cross-frequency phase–amplitude coupling (PAC) analysis at the Pz electrode, illustrating the relationship between low-frequency phase (horizontal axis) and higher-frequency amplitude (vertical axis). Coupling strength was estimated using the directPAC method and visualized as a two-dimensional frequency–frequency representation. **d, Machine-learning classification.** Receiver operating characteristic (ROC) curves illustrating pairwise classification of AD versus HC, AD versus FTD and HC versus FTD using ensemble learning, k-nearest neighbours (KNN), linear discriminant analysis (LDA) and logistic regression. Colours distinguish the three classification comparisons, with the corresponding area under the ROC curve (AUC) reported in each panel. These examples demonstrate the integration of statistical feature extraction, functional connectivity, cross-frequency coupling and multivariate classification within MLE EEG Toolbox. Classification performance is presented as an illustration of computational workflow functionality rather than evidence of independently validated diagnostic biomarkers.

